# Evaluation and Optimization of Chemically-Cleavable Linkers for Quantitative Mapping of Small Molecule-Protein Interactomes

**DOI:** 10.1101/654384

**Authors:** Adam J. Rabalski, Andrew R. Bogdan, Aleksandra Baranczak

**Affiliations:** Drug Discovery Science & Technology, AbbVie Inc., 1 N Waukegan Rd, North Chicago, Illinois 60064-6101 USA

## Abstract

Numerous reagents have been developed to enable chemical proteomic analysis of small molecule-protein interactomes. However, the performance of these reagents has not been systematically evaluated and compared. Herein, we report our efforts to conduct a parallel assessment of two widely-used chemically-cleavable linkers equipped with dialkoxydiphenylsilane (DADPS linker) and azobenzene (AZO linker) moieties. Profiling a cellular cysteinome using iodoacetamide alkyne probe demonstrated a significant discrepancy between the experimental results obtained through the application of each of the reagents. To better understand the source of observed discrepancy, a mass tolerant database search strategy using MSFragger software was performed. This resulted in identifying a previously unreported artifactual modification on the residual mass of the azobenzene linker. Furthermore, we conducted a comparative analysis of enrichment modes using both cleavable linkers. This effort determined that enrichment of proteolytic digests yielded a far greater number of identified cysteine residues than the enrichment conducted prior to protein digest. Inspired by recent studies where multiplexed quantitative labeling strategies were applied to cleavable biotin linkers, we combined this further optimized protocol using the DADPS cleavable linker with tandem mass tag (TMT) labeling to profile the FDA-approved covalent EGFR kinase inhibitor dacomitinib against the cysteinome of an epidermoid cancer cell line. Our analysis resulted in the detection and quantification of over 10,000 unique cysteine residues, a nearly 3-fold increase over previous studies that used cleavable biotin linkers for enrichment. Critically, cysteine residues corresponding to proteins directly as well as indirectly modulated by dacomitinib treatment were identified. Overall, our study suggests that the dialkoxydiphenylsilane linker could be broadly applied wherever chemically cleavable linkers are required for chemical proteomic characterization of cellular proteomes.

## Introduction

Chemical proteomics enables unbiased identification of small molecule-protein interactions in cellular systems^*1*^. One widely applied technique in chemical proteomics, termed activity-based protein profiling (ABPP), has been utilized in the identification of small molecule protein interactomes, characterization of entire protein families and probing the biology of proteins-of-interest^*2*^. More recently, the application of small molecule fragments to probe the reactivity of the cellular proteome has enabled new ways to target portions of the proteome long considered “undruggable”^*3–7*^. In most of the aforementioned studies, two critical questions must be addressed: the determination of the identities of the targeted proteins and the actual site of interaction between the small molecule and its protein target(s). Both questions have been successfully grappled with by utilizing a combination of small molecule probes forming irreversible conjugates with proteins and reagents that allow for selective capture and identification of protein targets by tandem mass spectrometry^*2*^.

Critically, a form of bioorthogonal transformation – copper catalyzed alkyne azide cycloaddition (CuAAC) – has been utilized to selectively couple small molecule probe-protein conjugates after cellular treatment with a tag that subsequently enables enrichment of engaged proteins and isolation of the probe-peptide adduct (Figure 1A). Those tags, typically referred to as cleavable biotin linkers, consist of three critical components: an azide group capable of reacting with an alkyne-derivatized fragment conjugated to a protein, a biotin group for enrichment, and finally, a cleavable moiety that allows for selective isolation of the adducted peptide (Figure 1B)^*8*^. Pioneering work by Speers et al. utilized an enzymatically-cleaved biotin linker which enabled cellular profiling of reactive sites in a proteome^*9, 10*^. This approach was further extended to profiling the reactive cysteinome in mouse and human cell extracts ^*11*^.

**Figure 1.**
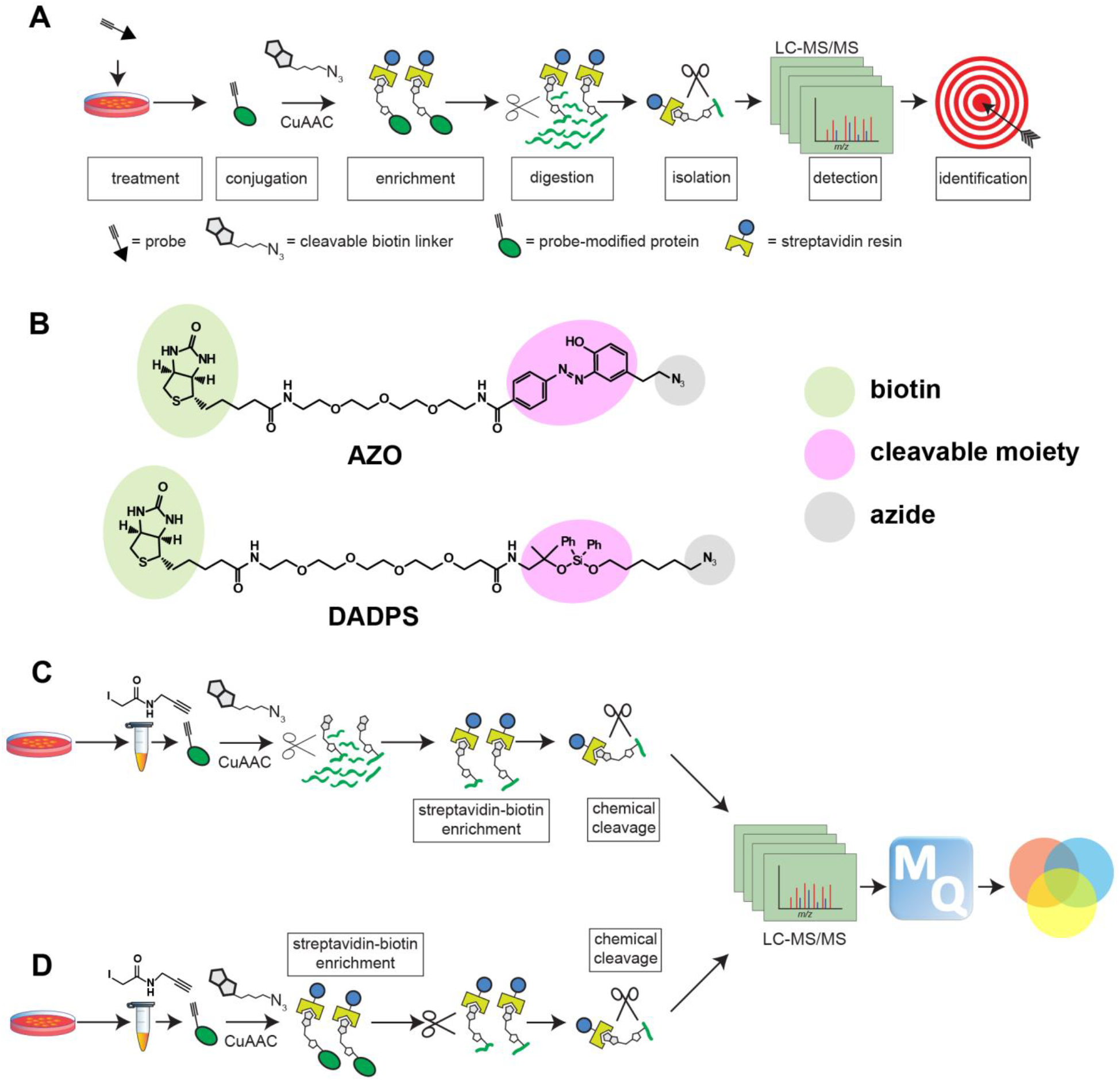
Representation of workflows and probes used in proteomic profiling of the reactive cysteinome with chemically-cleavable biotin linkers. A) General workflow for profiling reactive sites in the proteome. Samples are labeled with alkyne-derivatized probes then subjected to CuAAC with a cleavable biotin azide linker. Enrichment and digestion are followed by chemical cleavage, resulting in the identification of probe-modified peptides. B) Structures of azobenzene (AZO) and dialkoxydiphenylsilane (DADPS) cleavable biotin linkers used in the study C) Enrichment of cysteine residues post proteolytic digestion of proteins D) Enrichment of cysteine residues prior to proteolytic digestion of proteins.

More recently, modifications to cleavable biotin linkers have enabled a diverse set of chemically-cleavable reagents which provide the advantage of decreased sample processing times due to faster cleavage conditions and small residual modifications on conjugated peptides, a highly desirable property that can affect the ability to determine the identities of these peptides^*12, 13*^. Different chemical moieties have been implemented in the structure of cleavable linkers^*14–16*^. Among them, two types of linkers; azobenzene (AZO) linker and dialkoxydiphenylsilane (DADPS) linker, have been utilized most widely^*17, 18*^. The AZO linker has been applied in studies of the reactive cysteinome^*19, 20*^ and selenocysteinome^*21*^ in addition to the monitoring of newly synthesized bacterial proteins^*22*^. The biotin portion of the AZO linker is cleaved off upon reduction of the azobenzene group using sodium dithionite resulting in the formation of a residual aminophenol moiety on the cleaved peptide. To date, the applications of the DADPS linker included: identification of glycosylation sites^*23*^, profiling the deubiquitinase family of proteins^*24*^, and mapping the sites of small-molecule protein interactions^*25–27*^. The biotin portion of the DADPS linker is cleaved using formic acid leaving a hydroxyl moiety on the peptide adduct.

While each linker has been applied in numerous studies, so far there has been little understanding behind the criteria for their selection for a specific experimental application. Therefore, we decided to compare the performance of the AZO and DADPS cleavable linkers under similar experimental conditions. We were interested in understanding if any differences in the experimental results would arise through application of either reagent. To this end, we employed both linkers in an effort to profile the cellular cysteinome. Considering previous utilization of the AZO linker in characterizing the cysteinome, our goal was not to simply recapitulate reported work but to also investigate potential improvements to currently existing protocols. Moreover, in our studies, we took advantage of a mass tolerant search strategy with MSFragger^*28*^ software which allowed for the identification of unexpected modifications of the evaluated cleavable linkers. Finally, we demonstrated the utility of the developed methods in identifying the protein interactome of dacomitinib, a recently FDA-approved inhibitor of epidermal growth factor receptor (EGFR) kinase. This effort resulted in a better understanding of the selectivity profile of dacomitinib as well as the consequences of proteome modulation by this small molecule.

## Results and Discussion

In order to conduct a head-to-head comparison of the performance of the azobenzene (AZO) and dialkoxydiphenylsilane (DADPS) chemically-cleavable linkers in characterizing the cellular cysteinome (Figure 1B), we utilized the cysteine residue-reactive probe iodoacetamide alkyne (IAAyne) as it has been successfully deployed in chemical proteomic applications^*11, 19*^, and serves as a tool to profile cysteine-reactive covalent drugs used for disease indications^*29*^. We initiated our studies by subjecting 1 mg of lysate derived from K562 chronic myelogenous leukemia cells to alkylation in the presence of IAAyne (100 μM). This was followed by CuAAC-mediated derivatization of IAAyne-modified cysteines using either cleavable biotin azide. Subsequently, standard chloroform-methanol precipitation was used to remove excess CuAAC reagents.

Previous reports suggested that streptavidin-biotin enrichment conducted after proteolysis could result in a higher number of identifications than enrichment conducted prior to protein digestion^*30–32*^. We therefore decided to perform two enrichment approaches in parallel to understand how either method would affect the total number of identifications. For the first sample, biotin-mediated enrichment followed protein digestion while for the second sample the enrichment was performed prior to proteolysis (Figure 1C-D). Following enrichment and stringent washing, samples were subjected to chemical cleavage with sodium dithionite (AZO) or 10% formic acid (DADPS) to enable removal of the biotin moiety and isolation of the IAAyne-modified peptide adducts from the streptavidin resin. Samples were desalted using StageTips and LC-MS/MS analysis was conducted using a Thermo Orbitrap Fusion MS.

Label-free analysis using MaxQuant was performed to identify unique cysteine residues that could be isolated using either cleavable biotin linker. We analyzed the number of unique cysteine residues in each replicate and tabulated cysteine residues which had a localization probability greater than 0.80. Only localization scores from singly-modified cysteine containing peptides were used. Consequently, 11,400 and 10,316 unique cysteine residues were identified with the use of the DADPS linker when peptide and protein enrichment were performed, respectively. 7866 cysteine residues were identified in all three replicates when enrichment was performed post proteolysis, and 6676 cysteine residues were identified in all three replicates when enrichment was performed prior to proteolysis (Figure 2A). The application of the AZO linker enabled the identification of 9362 and 6664 total unique cysteine residues when performing peptide and protein enrichment, respectively. 4326 cysteine residues were identified in all three replicates when enrichment was performed after proteolysis, and 2795 cysteine residues were identified in all three replicates when enrichment was performed prior to proteolysis (Figure 2B). Overall, comparison of the four datasets demonstrated that the application of the DADPS biotin linker yielded the greatest number of unique cysteine residues when enrichment was performed after proteolytic digestion (Figure 2 & Table 1).

**Table 1.** Unique Cysteine Residues Identified by Chemically-Cleavable Linkers

| Linker | Enrichment Type | # Identifications in 3/3 replicates | Total # Identifications |
| --- | --- | --- | --- |
| DADPS | Protein | 6676 | 10316 |
| AZO | Protein | 2795 | 6664 |
| DADPS | Peptide | 7866 | 11440 |
| AZO | Peptide | 4326 | 9362 |

**Figure 2.**
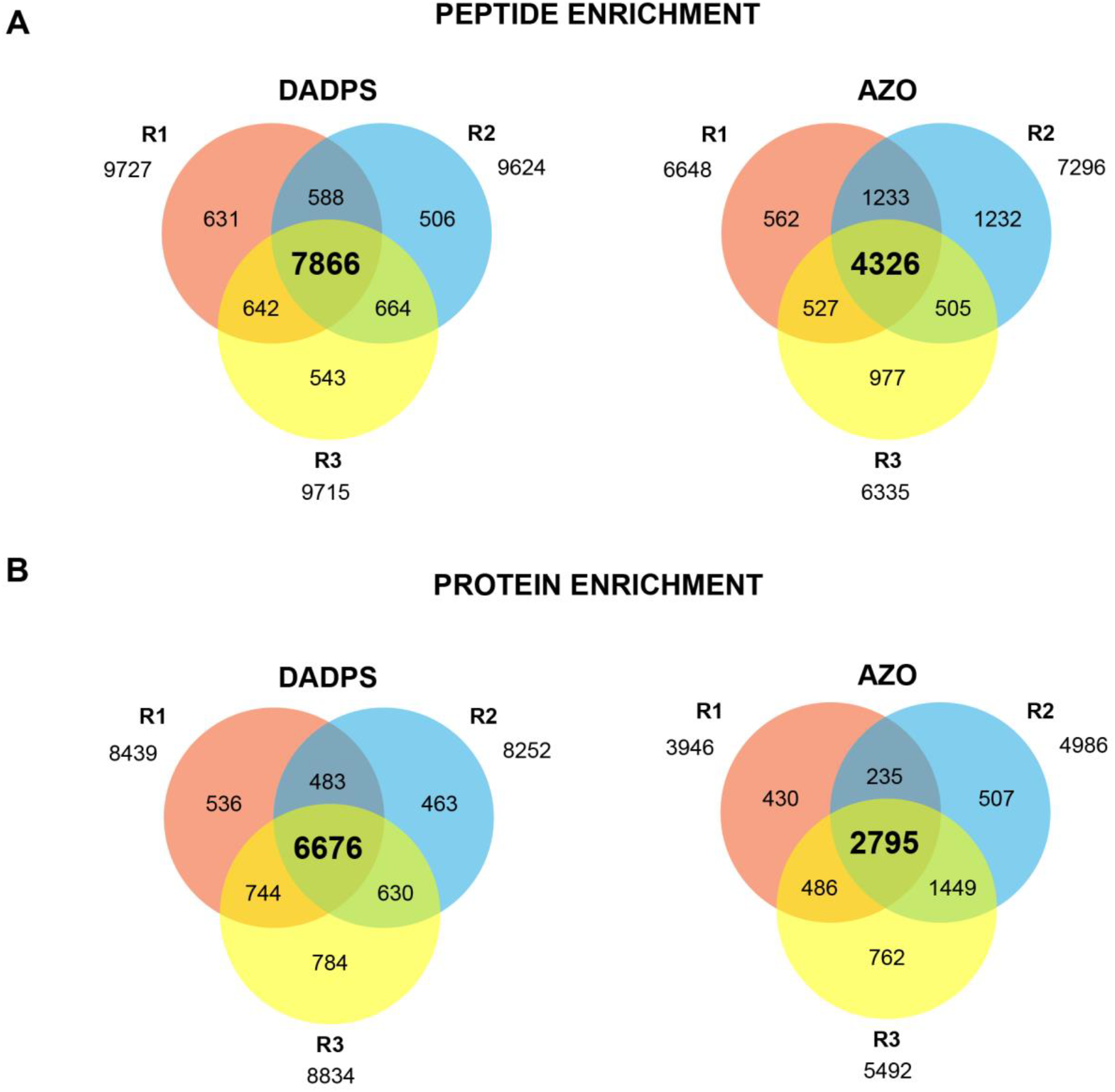
Comparison of the number of unique reactive cysteine residues detected by proteomics using DADPS and AZO linkers. All tabulated results included cysteine residues where localization probability was greater than 0.80 in at least one of the three replicates. A) Number of unique reactive cysteine residues identified when performing enrichment after proteolysis (n=3). B) Number of unique cysteine residues identified when performing enrichment before proteolysis (n =3).

A sizeable discrepancy was observed when the AZO linker was utilized indifferent of the enrichment method as each respective method yielded approximately twice as many unique identifications when the DADPS linker was applied. We decided to analyze the total number of proteins that would be identified from an on-bead digest when using either linker for enrichment of proteins prior to proteolytic digestion. Our expectation was that equivalent numbers of protein groups should be identified when controlling for the amount of material being labeled during the CuAAC reaction. Indeed, comparison of protein groups identified through the application of either linker did not reveal bias in enrichment of proteins, with either reagent capturing 90% of the 3502 protein groups identified (Figure 3A). In contrast, there was only a 54% overlap between the AZO and DADPS linkers when comparing identified cysteine residues in aggregate. This constituted 7612 cysteine residues out of a total of 14,098. 29.7% of the cysteines were exclusively identified when using the DADPS linker, while only 15% of the cysteine residues were exclusively identified when using AZO linker (Figure 3B).

**Figure 3.**
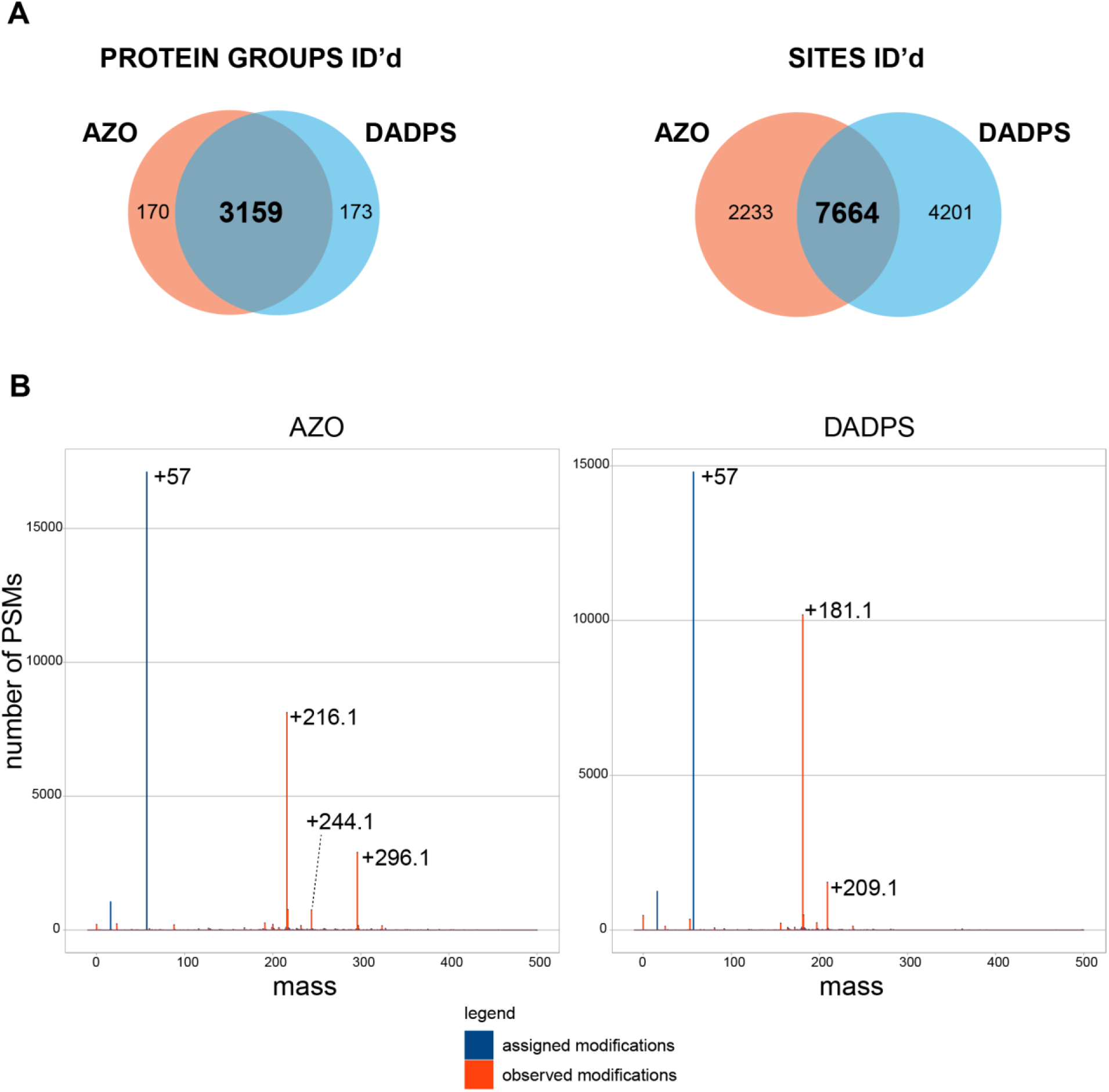
Analysis of yield discrepancies between AZO and DADPS linkers. A) Venn diagram of protein groups and reactive sites identified by either linker. Nearly identical number of protein groups were identified between data sets generated from on-bead digests through the application of the AZO linker and DADPS linker. In contrast, comparison of the total number of unique cysteine residues revealed a difference between the same data sets. B) MSFragger was used for a mass tolerant search to identify possible modifications missed in preliminary database searches. “Assigned modifications” represent modifications that were included as default during database searching (M oxidation +15.99490 Da and C carbamidomethylation +57.021464 Da) and “observed modifications” represent modifications that were not preset in database searches. Identification of a mass shift unique to the AZO linker (+80 Da) suggested possible artifactual modification of peptides when using this reagent.

We were interested in understanding whether the discrepancies between the number of identified cysteine residues could be attributed to an incorrect mass calculation for variable modifications during database searching or whether the presence of an additional uncharacterized modification resulted in a fewer number of identified unique cysteine residues when the AZO linker was utilized. To address these questions, we performed mass tolerant database searching with MSFragger^*28*^. Mass tolerant searching enables the identification of modifications that are not accounted for during a closed search with preset variable modifications and it has been demonstrated to increase the number of identifications from tandem mass spectra^*33*^. In order to attain high mass accuracy required for the mass tolerant database analysis, samples were re-acquired using MS2 scans detected by the Orbitrap. By performing the database search with a fixed modification of +57 Da for cysteine carbamidomethylation, we observed the expected mass differences of +216.1 Da and +181.1 Da for the AZO linker and DADPS linker respectively, which corresponded to the expected residual masses resulting from chemical cleavage (Figure 3B).

Furthermore, we concluded that the presence of additional peptide spectrum matches with an observed modification (+244.1 Da for AZO and +209.1 Da for DADPS) corresponding to a difference of approximately 28 Da from the expected residual modification mass of either cleavable linker could be attributed to the formylation of S/T/K residues or N-termini^*34, 35*^. Finally, a unique mass shift of +296.1 Da was only observed in peptide spectrum matches from peptides that were enriched using the AZO linker (Figure 3B). Site localization of the +296.1 Da modification using PTMProphet assigned the modification to cysteine residues, suggesting that the cleaved residual AZO linker modification was being additionally modified by a mass of +80 Da (Supp Fig 1A). Taking into account the reductive conditions using sodium dithionite for cleavage of the AZO linker, we considered sulfation as a potential source of the mass difference in detected peptides. Such a mass difference, identified in only one of the two datasets, pointed towards artifactual modification of the cleavable linker, reducing the likelihood that a biologically relevant modification such as phosphorylation (+79.96633 Da) was observed.

It has been previously demonstrated that sodium dithionite could undergo decomposition to thiosulfate in phosphate-buffered saline under low pH conditions^*36, 37*^. Presence of thiosulfate could on the other hand result in artifactual sulfation of peptides^*38*^. We hypothesized that the aminophenol moiety formed after cleavage of the AZO linker could be a likely target of sulfation. We reasoned that the addition of an acid to the eluent prior to peptide desalting, a well-established part of the sample preparation protocol^*19, 39*^, could be the likely cause of dithionite decomposition. We therefore decided to conduct additional analysis and included the AZO-SULFO modification as a variable modification during database searching. Consequently, searching the cleavable linker data with the addition of the AZO-SULFO modification using MaxQuant resulted in the identification of a total of 10,000 unique cysteine residues. We identified 3856 unique AZO-SULFO modified cysteine residues in all three replicates (Supp Fig 1B). A substantial overlap (32%) between AZO and AZO-SULFO modified cysteine residues was observed, suggesting that sequence-unique cysteine residues existed in either modification form in the acquired data set (Supp Fig 1B). We reasoned that this finding might explain the discrepancy between the total number of cysteine residues identified when comparing the DADPS linker and AZO linker.

To further interrogate the source of observed artifactual modification of peptides resulting from the application of the AZO linker, we conducted an experiment in which recombinant glutathione S-transferase omega 1 (GSTO1) protein, previously reported to possess a highly reactive cysteine residue in its active site ^*11*^, was incubated with IAAyne and subsequently subjected to CuAAC-mediated labeling with the AZO linker. This was followed by cleavage of the biotin portion with sodium dithionite. Prior to protein precipitation the obtained samples were subjected to incubation with or without the addition of formic acid in order to understand the potential role this reagent played in the occurrence of the AZO-SULFO modification. Interestingly, the AZO-SULFO modification on the peptide corresponding to the active site of GSTO1 was detected under both conditions (Supp Fig 2). Based on these results, we hypothesized that the potential decomposition of sodium dithionite occurred prior to the addition of formic acid. This could suggest that either reagents used for CuAAC labeling or sample processing for mass spectrometry might not be directly compatible with the application of sodium dithionite. However, this hypothesis would require further empirical validation.

Upon completion of our comparative study resulting in the identification of the DADPS linker as the more suitable reagent for chemical proteomic analysis of the cellular cysteinome, we decided to evaluate the applicability of the developed methods to characterize the protein-interactome of a drug-like molecule. We selected the covalent EGFR kinase inhibitor dacomitinib, which was recently approved for the treatment of non-small cell lung cancer^*40, 41*^. Dacomitinib is a pan-erythroblastic leukemia viral oncogene homologue (ERBB) inhibitor which targets wildtype and mutant forms of EGFR that possess activating L858R or exon 19 deletion mutations in addition to T790M mutation^*42, 43*^. For the purpose of our study, we utilized the A431 epidermoid cancer cell line as it was previously reported to be characterized by a high expression of wildtype EGFR^*44*^, and has been utilized in experiments profiling EGFR covalent inhibitors^*45, 46*^.

Previous reports indicated that the application of multiplexed labeling strategies for cleavable linkers could result in the identification of several thousand unique peptides within a single experiment when using reductive dimethyl and iTRAQ labeling strategies ^*47, 48*^. We therefore decided to implement a recently published protocol^*49*^ utilizing TMT labeling in combination with the DADPS linker (Supp Fig 3A). To this end, A431 epidermoid cancer cells were treated in the presence or absence of dacomitinib (5 μM) for 2 hours. Cells were harvested and cell extracts were subjected to labeling using a dacomitinib alkyne-derived probe. This was followed by CuAAC-mediated labeling to tag targets of the probe with a fluorophore (Supp Fig 3B). In-gel fluorescence analysis of treated samples as well as complementary immunoblot analysis using a phospho-specific antibody probing the activation status of EGFR confirmed on-target binding and successful inhibition of EGFR (Supp Fig 3C-D). Subsequently, cell lysates were incubated with IAAyne (100 μM) and derivatized with the DADPS linker. Proteomes were precipitated, washed, and proteolytically cleaved. TMT6plex labeling was performed using three replicate experiments and samples were analyzed using a Thermo Fusion Lumos MS. After data normalization and analysis (Supp Fig 4 & 5), a total of 10,201 unique cysteine residues were identified which represented a nearly 3-fold increase over recent studies in the number of unique cysteine residues observed when using IAAyne and a cleavable linker in a single experiment^*50*^. While the majority of the proteome and cysteinome were found to be unperturbed, a small proportion of quantified cysteine residues exceeded a magnitude log_2_ fold change of one which represented more than four standard deviations from the mean (Figure 4). We identified modulated cysteine residues that revealed increased or decreased labeling by the IAAyne probe in response to dacomitinib treatment. 15 cysteine residues corresponding to the sequence of EGFR did not demonstrate modulation, while the cysteine residue corresponding to the sequence of the active site of EGFR (C797) was not identified, likely due to the size of the peptide generated upon tryptic digestion.

**Figure 4.**
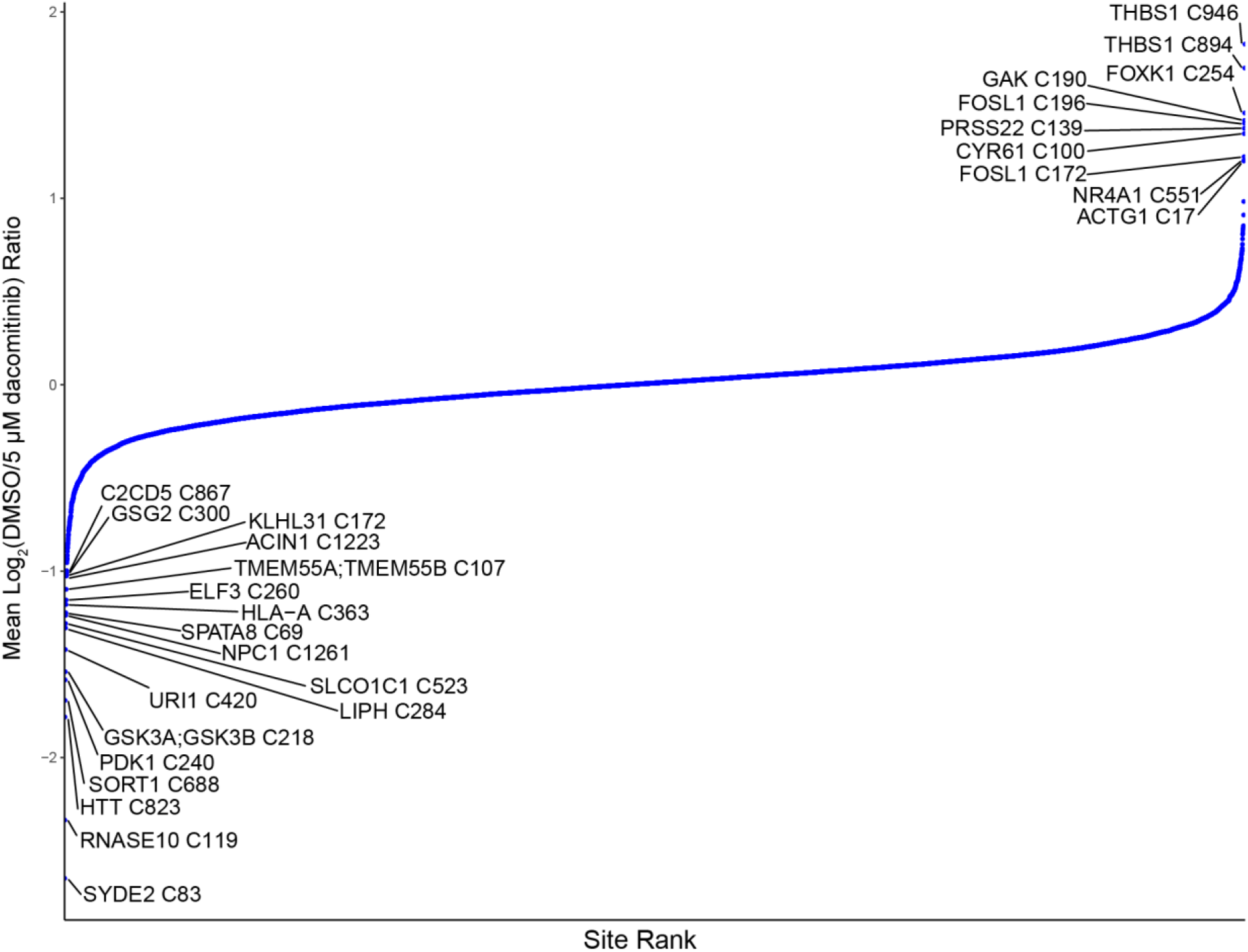
Characterization of the A431 cellular cysteinome upon dacomitinib treatment. Waterfall plot depicting cysteine residues identified using IAAyne in competition with the EGFR covalent inhibitor dacomitinib in cellular extracts derived from A431 cells. Cysteine residues that were competed or modulated to a greater extent than a magnitude log_2_ fold change of 1 are labeled with respective gene symbol and residue sequence position.

Interestingly, modulated cysteine residues occurred on several proteins where a single cysteine residue was perturbed relative to all other cysteine residues located on the respective protein (Supp Fig 6A). First, we identified C190 from Cyclin-G-associated kinase (GAK) that was engaged by IAAyne only in the absence of dacomitinib suggesting direct binding of dacomitinib to the kinase. This cysteine residue has been previously reported to be located next to the highly conserved DFG motif, which in an active kinase state coordinates magnesium ion in the ATP binding site^*51*^. Interestingly, a structurally similar analog of dacomitinib, FDA-approved EGFR inhibitor gefitinib, has also been demonstrated to target GAK^*29, 52, 53*^. Additionally, GAK C190 has been recently characterized as a target of a SM1-71, a promiscuous covalent kinase inhibitor ^*54*^.

Another cysteine residue that was observed to undergo modulation upon treatment with dacomitinib was C218, localized in close proximity to the active site of glycogen synthase kinase-3 beta (GSK3B). Interestingly, C218 was reported to be proximal to Y216 and phosphorylation of Y216 has been demonstrated to be required for GSK3B activity^*55*^. In our study, treatment with dacomitinib reduced phosphorylation of Y216 which was confirmed by immunoblot analysis (Supp Fig 6B). It has been previously reported that an electro-negative environment within the structure of a protein could destabilize the thiolate form and increase the pK_a_ of a cysteine^*56–58*^. In fact, the lack of change in protein abundance levels between dacomitinib treatment and control confirmed by immunoblot analysis (Supp Fig 6B) could suggest that upon reduction in GSK3B phosphorylation status, C218 becomes more nucleophilic and therefore more labeling of this cysteine residue occurred upon IAAyne treatment.

The increased labeling of a cysteine residue by IAAyne in response to dacomitinib treatment was also observed for another kinase, pyruvate dehydrogenase kinase 1 (PDK1). PDK1 C240 is proximal to a tyrosine phosphorylation site (Y243) which has been previously reported to be regulated by FGFR1 and is required for PDK1 activity ^*59*^. Previous proteomic studies have categorized this cysteine residue as hyper-reactive^*20*^. Moreover, it has been demonstrated that upon treatment with tyrosine kinase inhibitor TKI258, decreased phosphorylation of PDK1 Y243 results in decreased activity of PDK1^*59*^. The results of these reports combined with our proteomic analysis suggested that C240 reactivity could be governed by the phosphorylation status of the neighboring Y243 residue. As previously reported these findings may reflect potential cross-talk of cysteine reactivity with phosphorylation of proximal amino acid residues^*60, 61*^.

Instances of modulation where all cysteine residues identified from a protein were perturbed were also discovered. We identified three cysteine residues corresponding to the sequence of Fos-related antigen 1 (FOSL1), a transcription factor involved in the response to serum stimulation. It has been previously reported that upon serum stimulation, FOSL1 transcription increases and the protein is translated^*62*^. We confirmed by immunoblot that FOSL1 expression increased in the vehicle treatment and was down-regulated when cells were treated with dacomitinib (Supp Fig 6B). Our treatment conditions utilized fresh serum-containing media therefore the effect observed relative to the control treatment matched the effect of perturbation using dacomitinib. Interestingly, in a different example where a cysteine residue corresponding to the sequence of nuclear receptor subfamily 4 group A member 1 (NR4A1) was identified, changes in NR4A1 protein abundance were not observed despite the fact that NR4A1 has been previously implicated as a serum-responsive transcriptional regulator (Supp Fig 6B)^*63*^. While additional analysis would be required, these data suggested that NR4A1 C551 itself might be a direct target of dacomitinib and this interaction could play a role in interfering with serum-mediated transcriptional activation.

Overall, by taking into consideration the nature of our experiments, a general assessment of the type of identified cysteine residues allowed us to classify them into two categories. The first category included 10 out of 10,201 identified cysteine residues which were found to undergo a higher degree of labeling by IAAyne in the absence of dacomitinib. These cysteine residues could represent direct targets that were bound by dacomitinib or indirect targets which decreased in abundance in response to dacomitinib treatment. 18 cysteine residues comprised a second category, namely residues that were found to undergo a higher degree of labeling by IAAyne when cells were treated with dacomitinib relative to control treatment. These indirect effects may be due to increased protein abundance as a result of dacomitinib treatment or changes in cysteine reactivity due to differential regulation of post-translational modifications as a result of dacomitinib treatment.

In our study, we set out to compare two widely-used chemically-cleavable biotin linkers, previously applied in a variety of chemical proteomic studies. By conducting parallel profiling of the cellular cysteinome, we identified significant discrepancies between the number of identified cysteine residues enabled through the application of each of the linkers. Namely, we found that the application of the DADPS linker led to identification of a higher number of unique cysteine residues than the application of the AZO linker. Mass tolerant database searching using MSFragger enabled detection of an unexpected modification that was the possible source of the observed discrepancies. Specifically, we identified an artifactual modification in the form of sulfation on the azobenzene cleavable linker. This finding suggests that the utility of the AZO linker in chemical proteomic applications could be hindered as a substantial proportion of modification sites could become sulfated, potentially interfering with quantitative aspects of the assay or foregoing detection entirely. On the contrary, the compatibility of the cleavage conditions used to cleave the DADPS linker with mass spectrometry sample preparation protocols suits its application in profiling reactive sites in proteomes and makes it broadly applicable for characterizing small molecule protein interactomes.

## Methods

Detailed experimental methods can be found in the supporting information.

## Acknowledgements

We thank James P. Kath and Ryan A. McClure for critical reading of the manuscript.

## References

1. Drewes, G., and Knapp, S. (2018) Chemoproteomics and Chemical Probes for Target Discovery, Trends Biotechnol 36, 1275–1286.

2. Niphakis, M. J., and Cravatt, B. F. (2014) Enzyme inhibitor discovery by activity-based protein profiling, Annu Rev Biochem 83, 341–377.

3. Parker, C. G., Galmozzi, A., Wang, Y., Correia, B. E., Sasaki, K., Joslyn, C. M., Kim, A. S., Cavallaro, C. L., Lawrence, R. M., Johnson, S. R., Narvaiza, I., Saez, E., and Cravatt, B. F. (2017) Ligand and Target Discovery by Fragment-Based Screening in Human Cells, Cell 168, 527–541 e529.

4. Backus, K. M., Correia, B. E., Lum, K. M., Forli, S., Horning, B. D., Gonzalez-Paez, G. E., Chatterjee, S., Lanning, B. R., Teijaro, J. R., Olson, A. J., Wolan, D. W., and Cravatt, B. F. (2016) Proteome-wide covalent ligand discovery in native biological systems, Nature 534, 570–574.

5. Bar-Peled, L., Kemper, E. K., Suciu, R. M., Vinogradova, E. V., Backus, K. M., Horning, B. D., Paul, T. A., Ichu, T. A., Svensson, R. U., Olucha, J., Chang, M. W., Kok, B. P., Zhu, Z., Ihle, N. T., Dix, M. M., Jiang, P., Hayward, M. M., Saez, E., Shaw, R. J., and Cravatt, B. F. (2017) Chemical Proteomics Identifies Druggable Vulnerabilities in a Genetically Defined Cancer, Cell 171, 696–709 e623.

6. Hacker, S. M., Backus, K. M., Lazear, M. R., Forli, S., Correia, B. E., and Cravatt, B. F. (2017) Global profiling of lysine reactivity and ligandability in the human proteome, Nat Chem 9, 1181–1190.

7. Matthews, M. L., He, L., Horning, B. D., Olson, E. J., Correia, B. E., Yates, J. R., 3rd, Dawson, P. E., and Cravatt, B. F. (2017) Chemoproteomic profiling and discovery of protein electrophiles in human cells, Nat Chem 9, 234–243.

8. Yang, Y., Fonović, M., and Verhelst, S. H. L. (2017) Cleavable Linkers in Chemical Proteomics Applications, In Activity-Based Proteomics: Methods and Protocols (Overkleeft, H. S., and Florea, B. I., Eds.), pp 185–203, Springer New York, New York, NY.

9. Speers, A. E., and Cravatt, B. F. (2005) A tandem orthogonal proteolysis strategy for high-content chemical proteomics, Journal of the American Chemical Society 127, 10018–10019.

10. Weerapana, E., Speers, A. E., and Cravatt, B. F. (2007) Tandem orthogonal proteolysis-activity-based protein profiling (TOP-ABPP)--a general method for mapping sites of probe modification in proteomes, Nat Protoc 2, 1414–1425.

11. Weerapana, E., Wang, C., Simon, G. M., Richter, F., Khare, S., Dillon, M. B., Bachovchin, D. A., Mowen, K., Baker, D., and Cravatt, B. F. (2010) Quantitative reactivity profiling predicts functional cysteines in proteomes, Nature 468, 790–795.

12. Ficarro, S. B., Browne, C. M., Card, J. D., Alexander, W. M., Zhang, T., Park, E., McNally, R., Dhe-Paganon, S., Seo, H. S., Lamberto, I., Eck, M. J., Buhrlage, S. J., Gray, N. S., and Marto, J. A. (2016) Leveraging Gas-Phase Fragmentation Pathways for Improved Identification and Selective Detection of Targets Modified by Covalent Probes, Anal Chem 88, 12248–12254.

13. Flaxman, H. A., and Woo, C. M. (2018) Mapping the Small Molecule Interactome by Mass Spectrometry, Biochemistry 57, 186–193.

14. Yang, Y., Hahne, H., Kuster, B., and Verhelst, S. H. (2013) A simple and effective cleavable linker for chemical proteomics applications, Mol Cell Proteomics 12, 237–244.

15. Nessen, M. A., Kramer, G., Back, J., Baskin, J. M., Smeenk, L. E., de Koning, L. J., van Maarseveen, J. H., de Jong, L., Bertozzi, C. R., Hiemstra, H., and de Koster, C. G. (2009) Selective enrichment of azide-containing peptides from complex mixtures, J Proteome Res 8, 3702–3711.

16. Kim, H. Y., Tallman, K. A., Liebler, D. C., and Porter, N. A. (2009) An azido-biotin reagent for use in the isolation of protein adducts of lipid-derived electrophiles by streptavidin catch and photorelease, Mol Cell Proteomics 8, 2080–2089.

17. Verhelst, S. H., Fonovic, M., and Bogyo, M. (2007) A mild chemically cleavable linker system for functional proteomic applications, Angew Chem Int Ed Engl 46, 1284–1286.

18. Szychowski, J., Mahdavi, A., Hodas, J. J., Bagert, J. D., Ngo, J. T., Landgraf, P., Dieterich, D. C., Schuman, E. M., and Tirrell, D. A. (2010) Cleavable biotin probes for labeling of biomolecules via azide-alkyne cycloaddition, Journal of the American Chemical Society 132, 18351–18360.

19. Qian, Y., Martell, J., Pace, N. J., Ballard, T. E., Johnson, D. S., and Weerapana, E. (2013) An isotopically tagged azobenzene-based cleavable linker for quantitative proteomics, Chembiochem 14, 1410–1414.

20. Bak, D. W., Pizzagalli, M. D., and Weerapana, E. (2017) Identifying Functional Cysteine Residues in the Mitochondria, ACS Chem Biol 12, 947–957.

21. Bak, D. W., Gao, J., Wang, C., and Weerapana, E. (2018) A Quantitative Chemoproteomic Platform to Monitor Selenocysteine Reactivity within a Complex Proteome, Cell Chem Biol 25, 1157–1167 e1154.

22. Yang, Y. Y., Grammel, M., Raghavan, A. S., Charron, G., and Hang, H. C. (2010) Comparative analysis of cleavable azobenzene-based affinity tags for bioorthogonal chemical proteomics, Chem Biol 17, 1212–1222.

23. Woo, C. M., Iavarone, A. T., Spiciarich, D. R., Palaniappan, K. K., and Bertozzi, C. R. (2015) Isotope-targeted glycoproteomics (IsoTaG): a mass-independent platform for intact N- and O-glycopeptide discovery and analysis, Nat Methods 12, 561–567.

24. Hewings, D. S., Flygare, J. A., Bogyo, M., and Wertz, I. E. (2017) Activity-based probes for the ubiquitin conjugation-deconjugation machinery: new chemistries, new tools, and new insights, FEBS J 284, 1555–1576.

25. Wang, J., Zhang, C. J., Zhang, J., He, Y., Lee, Y. M., Chen, S., Lim, T. K., Ng, S., Shen, H. M., and Lin, Q. (2015) Mapping sites of aspirin-induced acetylations in live cells by quantitative acid-cleavable activity-based protein profiling (QA-ABPP), Sci Rep 5, 7896.

26. Li, W., Zhou, Y., Tang, G., and Xiao, Y. (2016) Characterization of the Artemisinin Binding Site for Translationally Controlled Tumor Protein (TCTP) by Bioorthogonal Click Chemistry, Bioconjug Chem 27, 2828–2833.

27. Gao, J., Mfuh, A., Amako, Y., and Woo, C. M. (2018) Small Molecule Interactome Mapping by Photoaffinity Labeling Reveals Binding Site Hotspots for the NSAIDs, Journal of the American Chemical Society.

28. Kong, A. T., Leprevost, F. V., Avtonomov, D. M., Mellacheruvu, D., and Nesvizhskii, A. I. (2017) MSFragger: ultrafast and comprehensive peptide identification in mass spectrometry-based proteomics, Nat Methods 14, 513–520.

29. Chaikuad, A., Koch, P., Laufer, S. A., and Knapp, S. (2018) The Cysteinome of Protein Kinases as a Target in Drug Development, Angewandte Chemie International Edition 57, 4372–4385.

30. Schiapparelli, L. M., McClatchy, D. B., Liu, H. H., Sharma, P., Yates, J. R., 3rd, and Cline, H. T. (2014) Direct detection of biotinylated proteins by mass spectrometry, J Proteome Res 13, 3966–3978.

31. Yang, J., Gupta, V., Tallman, K. A., Porter, N. A., Carroll, K. S., and Liebler, D. C. (2015) Global, in situ, site-specific analysis of protein S-sulfenylation, Nat Protoc 10, 1022–1037.

32. Qin, K., Zhu, Y., Qin, W., Gao, J., Shao, X., Wang, Y. L., Zhou, W., Wang, C., and Chen, X. (2018) Quantitative Profiling of Protein O-GlcNAcylation Sites by an Isotope-Tagged Cleavable Linker, ACS Chem Biol 13, 1983–1989.

33. Chick, J. M., Kolippakkam, D., Nusinow, D. P., Zhai, B., Rad, R., Huttlin, E. L., and Gygi, S. P. (2015) A mass-tolerant database search identifies a large proportion of unassigned spectra in shotgun proteomics as modified peptides, Nat Biotechnol 33, 743–749.

34. Zheng, S., and Doucette, A. A. (2016) Preventing N- and O-formylation of proteins when incubated in concentrated formic acid, Proteomics 16, 1059–1068.

35. Devabhaktuni, A., Lin, S., Zhang, L., Swaminathan, K., Gonzalez, C. G., Olsson, N., Pearlman, S. M., Rawson, K., and Elias, J. E. (2019) TagGraph reveals vast protein modification landscapes from large tandem mass spectrometry datasets, Nature Biotechnology.

36. Danehy, J. P., and Zubritsky, C. W. (1974) Iodometric method for the determination of dithionite, bisulfite, and thiosulfate in the presence of each other and its use in following the decomposition of aqueous solutions of sodium dithionite, Anal Chem 46, 391–395.

37. Spencer, M. S. (1967) Chemistry of sodium dithionite. Part 1.—Kinetics of decomposition in aqueous bisulphite solutions, Trans. Faraday Soc. 63, 2510–2515.

38. Gharib, M., Marcantonio, M., Lehmann, S. G., Courcelles, M., Meloche, S., Verreault, A., and Thibault, P. (2009) Artifactual sulfation of silver-stained proteins: implications for the assignment of phosphorylation and sulfation sites, Mol Cell Proteomics 8, 506–518.

39. Abo, M., Li, C., and Weerapana, E. (2018) Isotopically-Labeled Iodoacetamide-Alkyne Probes for Quantitative Cysteine-Reactivity Profiling, Mol Pharm 15, 743–749.

40. Wu, Y.-L., Cheng, Y., Zhou, X., Lee, K. H., Nakagawa, K., Niho, S., Tsuji, F., Linke, R., Rosell, R., Corral, J., Migliorino, M. R., Pluzanski, A., Sbar, E. I., Wang, T., White, J. L., Nadanaciva, S., Sandin, R., and Mok, T. S. (2017) Dacomitinib versus gefitinib as first-line treatment for patients with EGFR - mutation-positive non-small-cell lung cancer (ARCHER 1050): a randomised, open-label, phase 3 trial, The Lancet Oncology 18, 1454–1466.

41. Urquhart, L. (2018) Regulatory watch: FDA new drug approvals in Q3 2018. Nat Rev Drug Discov 17, 779.

42. Engelman, J. A., Zejnullahu, K., Gale, C. M., Lifshits, E., Gonzales, A. J., Shimamura, T., Zhao, F., Vincent, P. W., Naumov, G. N., Bradner, J. E., Althaus, I. W., Gandhi, L., Shapiro, G. I., Nelson, J. M., Heymach, J. V., Meyerson, M., Wong, K. K., and Janne, P. A. (2007) PF00299804, an irreversible pan-ERBB inhibitor, is effective in lung cancer models with EGFR and ERBB2 mutations that are resistant to gefitinib, Cancer Res 67, 11924–11932.

43. Gonzales, A. J., Hook, K. E., Althaus, I. W., Ellis, P. A., Trachet, E., Delaney, A. M., Harvey, P. J., Ellis, T. A., Amato, D. M., Nelson, J. M., Fry, D. W., Zhu, T., Loi, C. M., Fakhoury, S. A., Schlosser, K. M., Sexton, K. E., Winters, R. T., Reed, J. E., Bridges, A. J., Lettiere, D. J., Baker, D. A., Yang, J., Lee, H. T., Tecle, H., and Vincent, P. W. (2008) Antitumor activity and pharmacokinetic properties of PF-00299804, a second-generation irreversible pan-erbB receptor tyrosine kinase inhibitor, Mol Cancer Ther 7, 1880–1889.

44. Stanton, P., Richards, S., Reeves, J., Nikolic, M., Edington, K., Clark, L., Robertson, G., Souter, D., Mitchell, R., Hendler, F. J., and et al. (1994) Epidermal growth factor receptor expression by human squamous cell carcinomas of the head and neck, cell lines and xenografts, Br J Cancer 70, 427–433.

45. Lanning, B. R., Whitby, L. R., Dix, M. M., Douhan, J., Gilbert, A. M., Hett, E. C., Johnson, T. O., Joslyn, C., Kath, J. C., Niessen, S., Roberts, L. R., Schnute, M. E., Wang, C., Hulce, J. J., Wei, B., Whiteley, L. O., Hayward, M. M., and Cravatt, B. F. (2014) A road map to evaluate the proteome-wide selectivity of covalent kinase inhibitors, Nat Chem Biol 10, 760–767.

46. Niessen, S., Dix, M. M., Barbas, S., Potter, Z. E., Lu, S., Brodsky, O., Planken, S., Behenna, D., Almaden, C., Gajiwala, K. S., Ryan, K., Ferre, R., Lazear, M. R., Hayward, M. M., Kath, J. C., and Cravatt, B. F. (2017) Proteome-wide Map of Targets of T790M-EGFR-Directed Covalent Inhibitors, Cell Chem Biol 24, 1388–1400 e1387.

47. Yang, F., Gao, J., Che, J., Jia, G., and Wang, C. (2018) A Dimethyl-Labeling-Based Strategy for Site-Specifically Quantitative Chemical Proteomics, Anal Chem 90, 9576–9582.

48. Tian, C., Sun, R., Liu, K., Fu, L., Liu, X., Zhou, W., Yang, Y., and Yang, J. (2017) Multiplexed Thiol Reactivity Profiling for Target Discovery of Electrophilic Natural Products, Cell Chem Biol 24, 1416–1427 e1415.

49. Navarrete-Perea, J., Yu, Q., Gygi, S. P., and Paulo, J. A. (2018) Streamlined Tandem Mass Tag (SL-TMT) Protocol: An Efficient Strategy for Quantitative (Phospho)proteome Profiling Using Tandem Mass Tag-Synchronous Precursor Selection-MS3, J Proteome Res 17, 2226–2236.

50. Zaro, B. W., Vinogradova, E. V., Lazar, D. C., Blewett, M. M., Suciu, R. M., Takaya, J., Studer, S., de la Torre, J. C., Casanova, J.-L., Cravatt, B. F., and Teijaro, J. R. (2019) Dimethyl Fumarate Disrupts Human Innate Immune Signaling by Targeting the IRAK4–MyD88 Complex, The Journal of Immunology, ji1801627.

51. Hanks, S. K., and Hunter, T. (1995) Protein kinases 6. The eukaryotic protein kinase superfamily: kinase (catalytic) domain structure and classification, The FASEB Journal 9, 576–596.

52. Brehmer, D., Greff, Z., Godl, K., Blencke, S., Kurtenbach, A., Weber, M., Muller, S., Klebl, B., Cotten, M., Keri, G., Wissing, J., and Daub, H. (2005) Cellular targets of gefitinib, Cancer Res 65, 379–382.

53. Sharma, K., Weber, C., Bairlein, M., Greff, Z., Keri, G., Cox, J., Olsen, J. V., and Daub, H. (2009) Proteomics strategy for quantitative protein interaction profiling in cell extracts, Nat Methods 6, 741–744.

54. Rao, S., Gurbani, D., Du, G., Everley, R. A., Browne, C. M., Chaikuad, A., Li, T., Schroder, M., Gondi, S., Ficarro, S. B., Sim, T., Kim, N. D., Berberich, M. J., Knapp, S., Marto, J. A., Westover, K. D., Sorger, P. K., and Gray, N. S. (2019) Leveraging Compound Promiscuity to Identify Targetable Cysteines within the Kinome, Cell Chem Biol.

55. Hughes, K., Nikolakaki, E., Plyte, S. E., Totty, N. F., and Woodgett, J. R. (1993) Modulation of the glycogen synthase kinase-3 family by tyrosine phosphorylation, EMBO J 12, 803–808.

56. Harris, T. K., and Turner, G. J. (2002) Structural basis of perturbed pKa values of catalytic groups in enzyme active sites, IUBMB Life 53, 85–98.

57. Salsbury, F. R., Jr., Knutson, S. T., Poole, L. B., and Fetrow, J. S. (2008) Functional site profiling and electrostatic analysis of cysteines modifiable to cysteine sulfenic acid, Protein Sci 17, 299–312.

58. Witt, A. C., Lakshminarasimhan, M., Remington, B. C., Hasim, S., Pozharski, E., and Wilson, M. A. (2008) Cysteine pKa depression by a protonated glutamic acid in human DJ-1, Biochemistry 47, 7430–7440.

59. Hitosugi, T., Fan, J., Chung, T. W., Lythgoe, K., Wang, X., Xie, J., Ge, Q., Gu, T. L., Polakiewicz, R. D., Roesel, J. L., Chen, G. Z., Boggon, T. J., Lonial, S., Fu, H., Khuri, F. R., Kang, S., and Chen, J. (2011) Tyrosine phosphorylation of mitochondrial pyruvate dehydrogenase kinase 1 is important for cancer metabolism, Mol Cell 44, 864–877.

60. Dong, M., Bian, Y., Dong, J., Wang, K., Liu, Z., Qin, H., Ye, M., and Zou, H. (2015) Selective Enrichment of Cysteine-Containing Phosphopeptides for Subphosphoproteome Analysis, J Proteome Res 14, 5341–5347.

61. Huang, H., Haar Petersen, M., Ibanez-Vea, M., Lassen, P. S., Larsen, M. R., and Palmisano, G. (2016) Simultaneous Enrichment of Cysteine-containing Peptides and Phosphopeptides Using a Cysteine-specific Phosphonate Adaptable Tag (CysPAT) in Combination with titanium dioxide (TiO2) Chromatography, Mol Cell Proteomics 15, 3282–3296.

62. Zippo, A., Serafini, R., Rocchigiani, M., Pennacchini, S., Krepelova, A., and Oliviero, S. (2009) Histone crosstalk between H3S10ph and H4K16ac generates a histone code that mediates transcription elongation, Cell 138, 1122–1136.

63. Nakai, A., Kartha, S., Sakurai, A., Toback, F. G., and DeGroot, L. J. (1990) A human early response gene homologous to murine nur77 and rat NGFI-B, and related to the nuclear receptor superfamily, Mol Endocrinol 4, 1438–1443.

