## Supplemental information for "Evaluation and Optimization of Chemically-Cleavable Linkers for Quantitative Mapping of Small Molecule-Protein Interactomes"

### Supplemental Figures

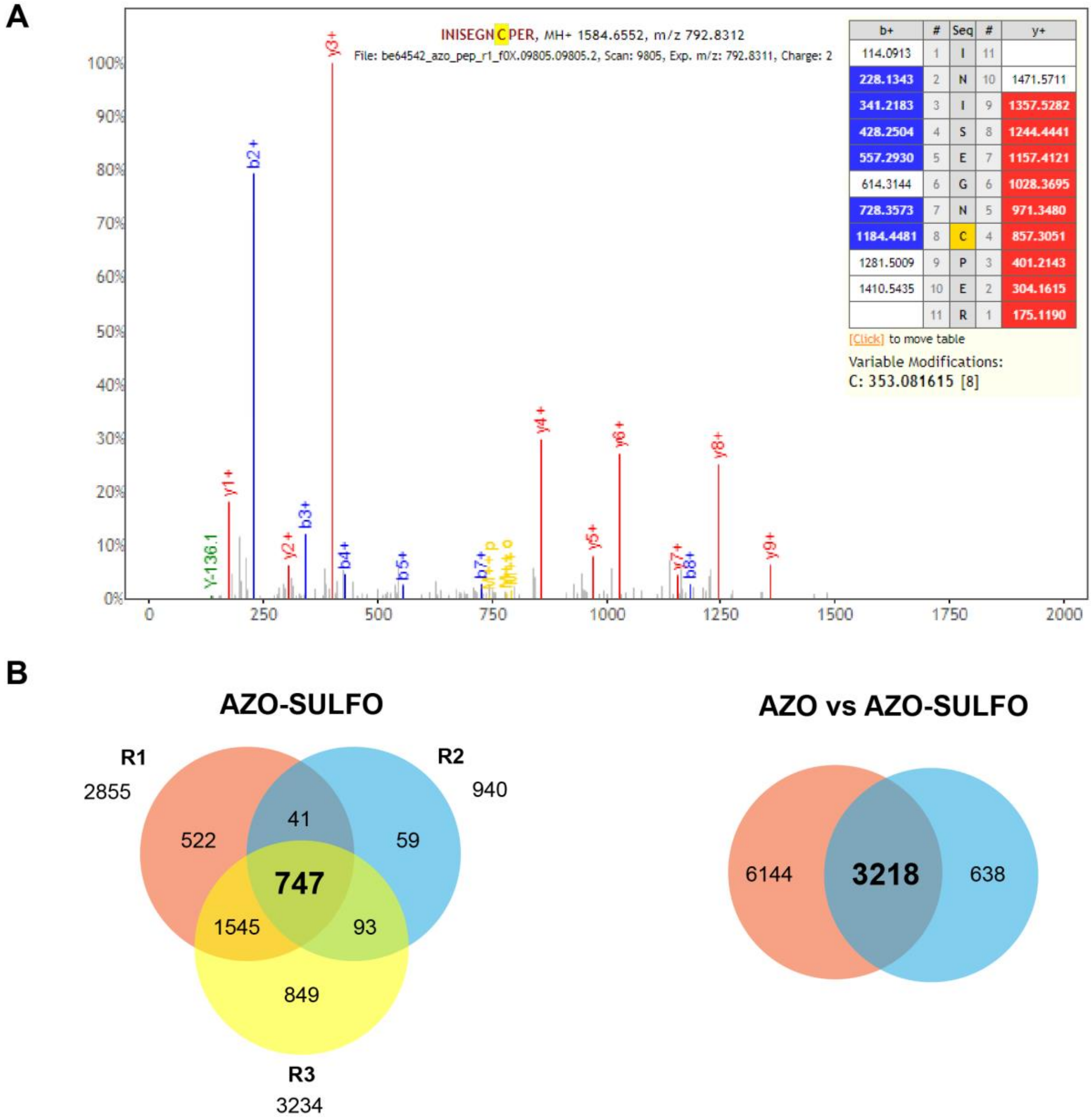

**Supplementary Figure 1. Identification of an artifactual modification on the AZO linker.** A) Mass spectrum showing the localization of sulfated azobenzene residual linker on a peptide from PCBP1 protein B) Comparison of cysteine residues modified by AZO-residual linker and sulfated-AZO-residual linker revealed substantial overlap between the two datasets.

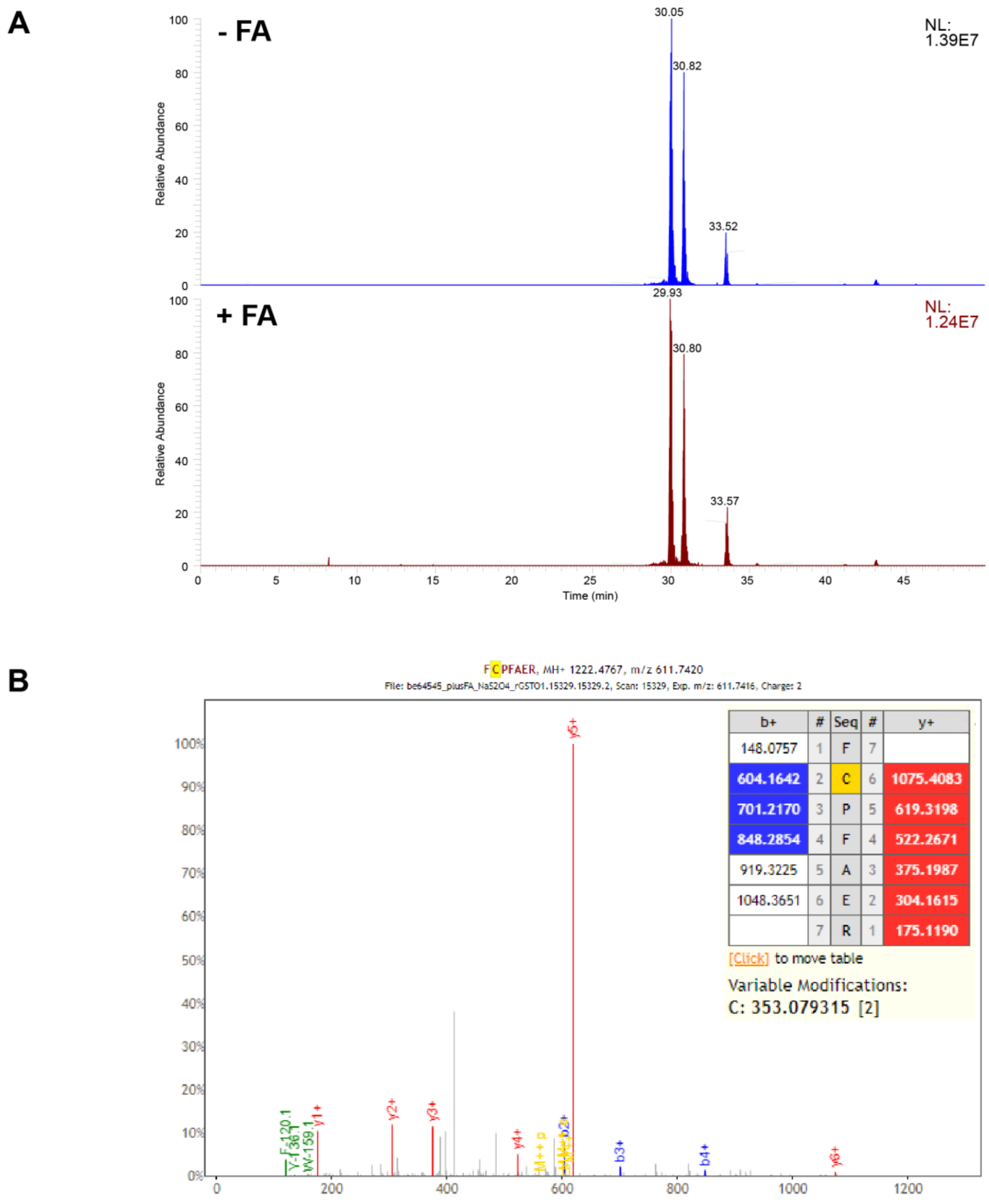

**Supplementary Figure 2. Evaluation of the AZO linker CuAAC protocol with the use of rGSTO1.** A) rGSTO1 was labeled with IAAyne and AZO biotin linker, and subjected to sodium dithionite-mediated cleavage in the presence (+FA) or absence (-FA) of formic acid. Extracted ion chromatograms of rGSTO1 active site peptide FCPFAER showed equal amount of sulfation suggesting that decomposition of sodium dithionite occurred prior to the addition of formic acid. B) Mass spectrum of sulfated-AZO derivatized peptide (FCPFAER) corresponding to the active site of rGSTO1.

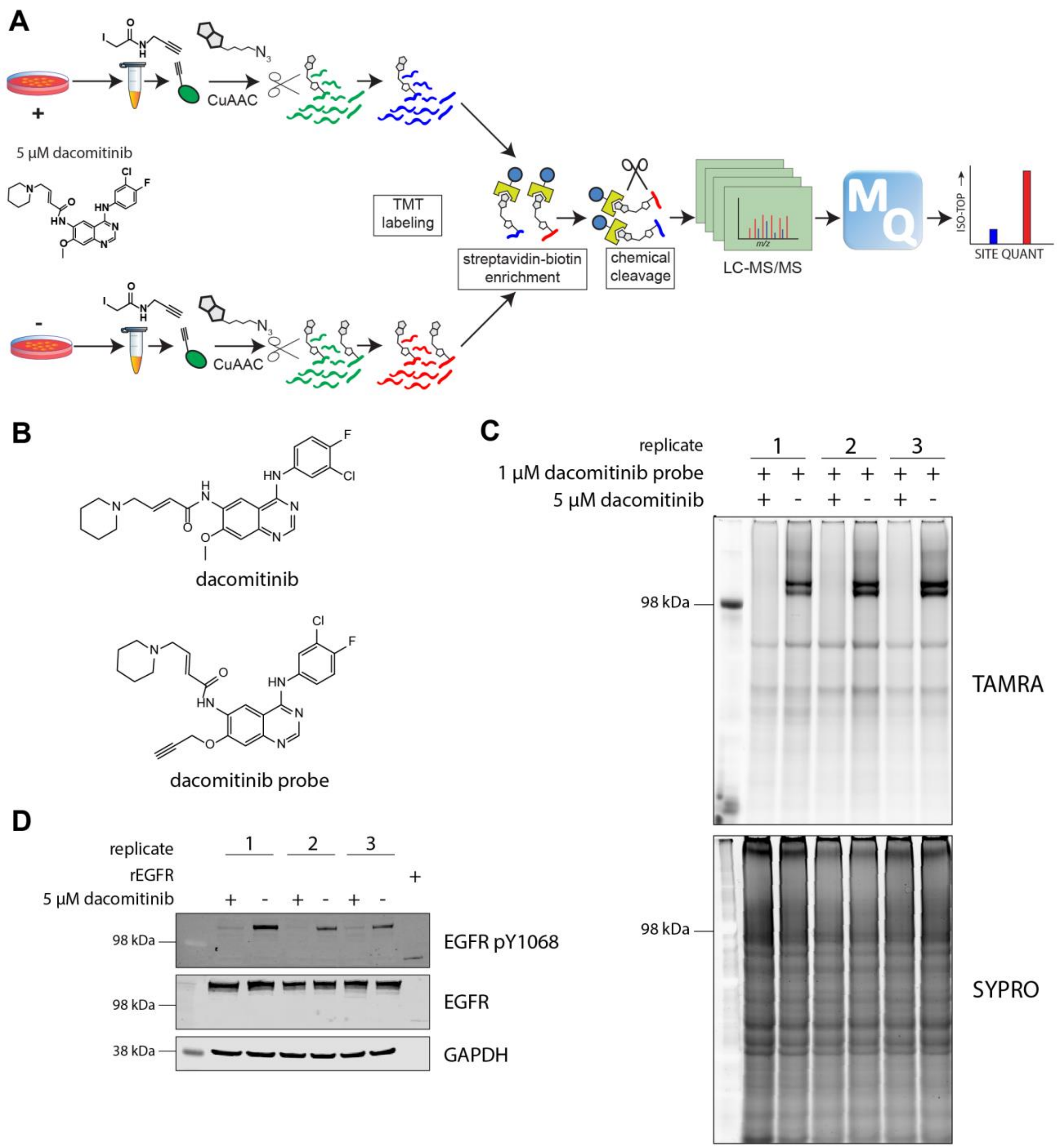

**Supplementary Figure 3. Application of chemically-cleavable DAPDS linker for identification of the protein interactome of the covalent EGFR kinase inhibitor dacomitinib.** A) Schematic representation of TMT workflow used for profiling of the proteome derived from A431 cells treated with dacomitinib. B) Structures of dacomitinib and alkyne-derivatized dacomitinib probe. C) In-gel fluorescence-based analysis of cellular extracts derived from A431 cells treated with dacomitinib. Cells were subjected to incubation in the presence or absence of dacomitinib in serum-containing media for 2 hours. Cell extracts were incubated with an alkyne-derivatized dacomitinib probe and subjected to CuAAC labeling with TAMRA fluorophore. Labeled extracts were separated using SDS-PAGE and visualized using a fluorescence scanner. D) Immunoblot-based analysis of cellular extracts derived from A431 cells treated with dacomitinib. PVDF membranes were stained with phospho-specific Y1068 EGFR antibody to reveal the effect of dacomitinib treatment. GAPDH was used as a loading control. The analyses confirmed direct target engagement of EGFR by dacomitinib.

A

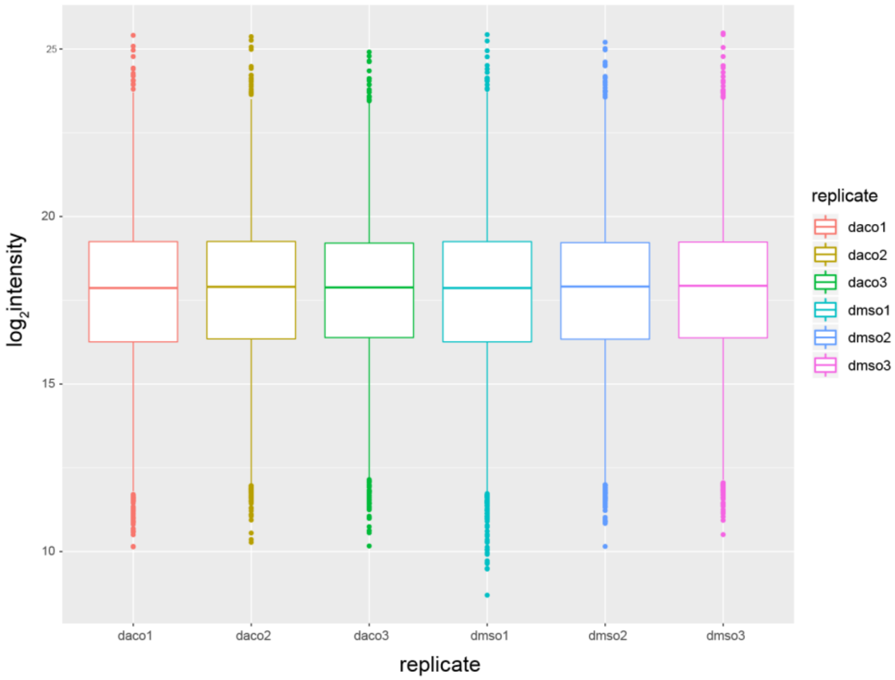

B

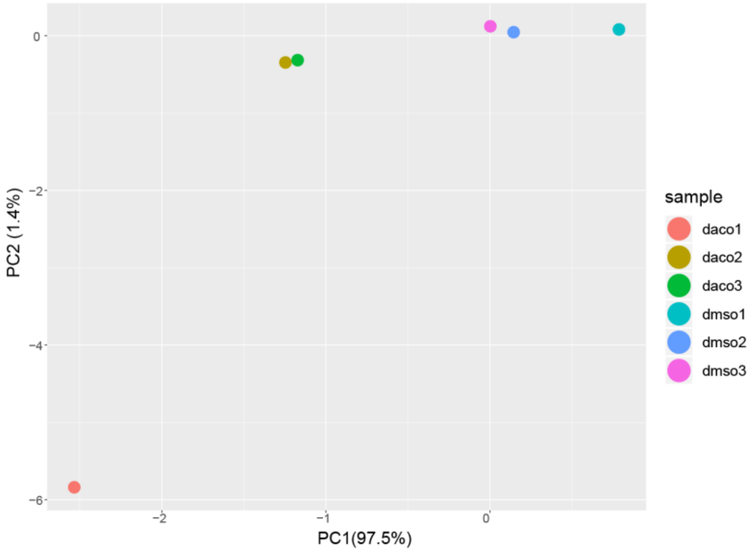

**Supplementary Figure 4. Comparisons of TMT intensities for cysteine residues across replicates and conditions.** A) Boxplot demonstrating $log_2$ intensities of proteome-wide cysteine residues from replicates treated with dacomitinib or vehicle following normalization. B) PCA analysis of $log_2$ intensities of modified cysteine residues showed separation between dacomitinib treated and vehicle treated samples.

A

B

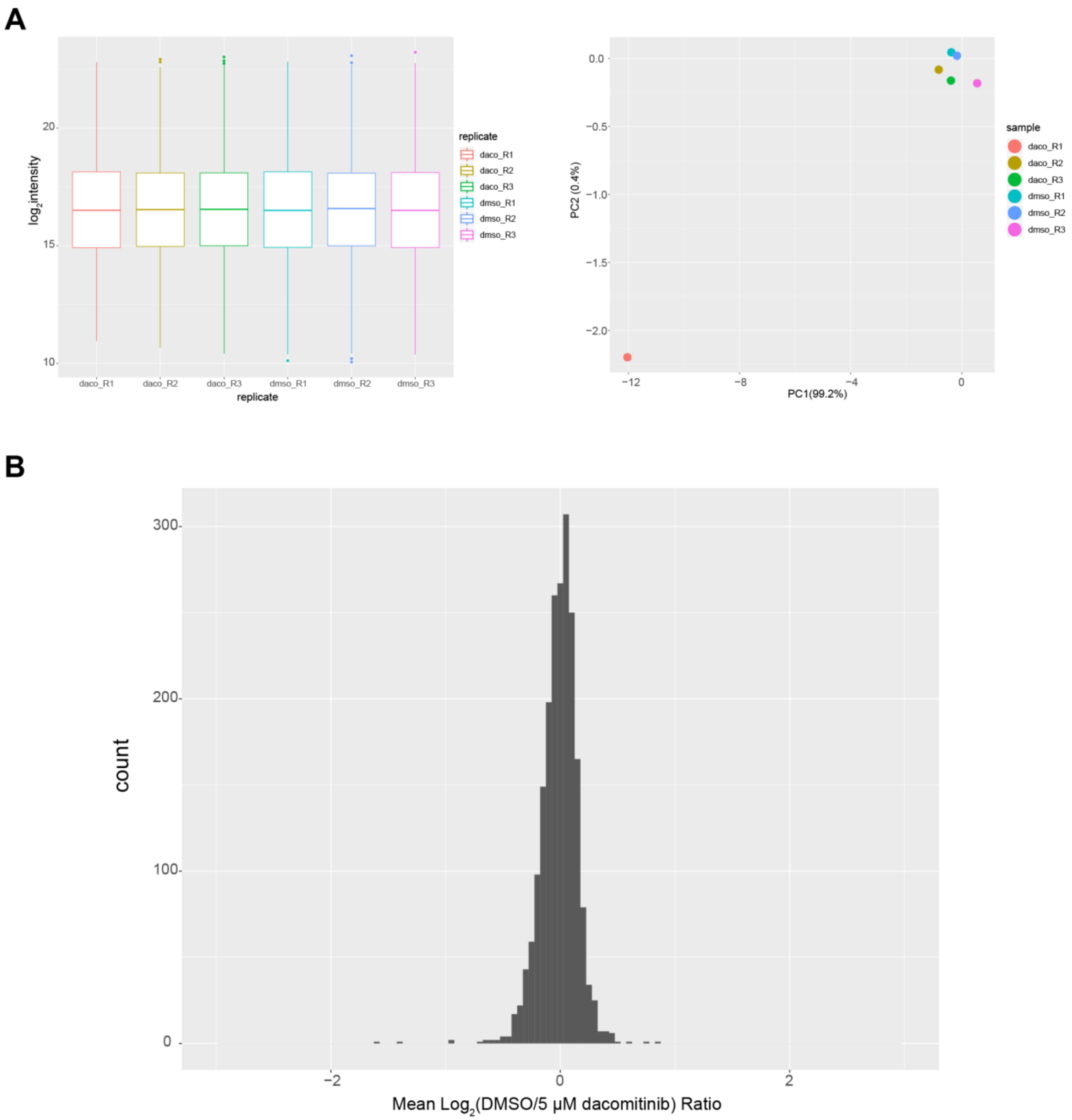

**Supplementary Figure 5. Comparisons of TMT intensities for proteins across replicates and conditions.** A) Boxplot demonstrating $log_2$ intensities of all detected proteins from replicates treated with dacomitinib or vehicle following normalization. B) Histogram of mean $log_2$ ratios of protein abundance between dacomitinib treatment and control demonstrated proteome integrity during 2 hour treatment with dacomitinib. The majority of protein abundances did not change and majority of protein abundance ratios observed were smaller in magnitude than observed cysteine residue ratios (See Figure 4 and Supp Fig 6).

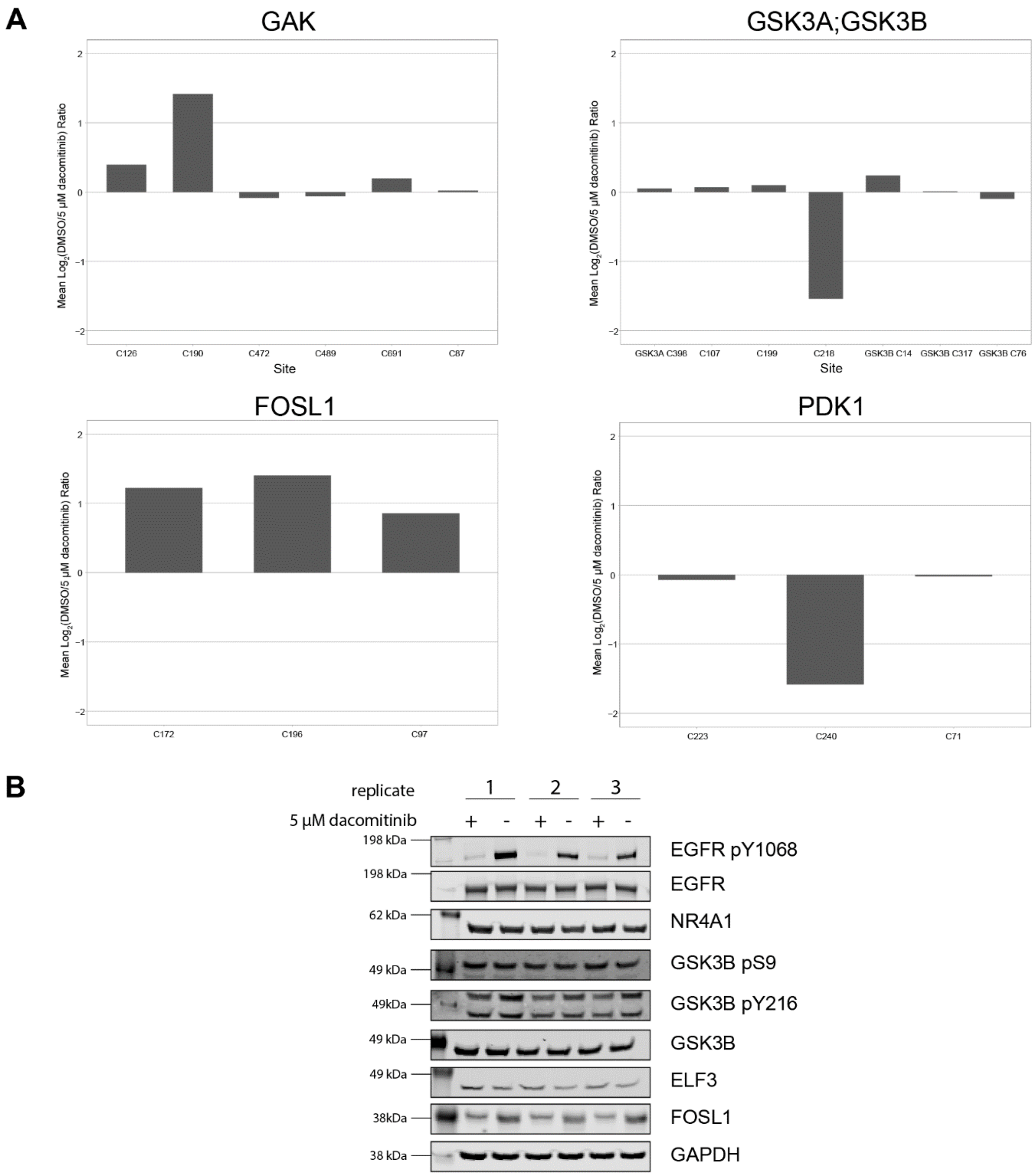

**Supplementary Figure 6. Analysis of cysteine residues modulated in response to dacomitinib perturbation.** A) Barplots depicting quantitative response upon dacomitinib treatment. Modulation of singular cysteine residues (independent of other cysteines on the same protein) suggested direct or indirect modulation as a result of dacomitinib treatment and an effect that was independent of a change in protein abundance. Modulation of several cysteine residues with the same magnitude change indicated quantitation that was dependent on protein abundance changes. B) Immunoblot analysis corroborated results observed from chemical proteomics data. Cellular extracts derived from A431 cells treated with dacomitinib were separated by SDS-PAGE and transferred to PVDF membranes. Membranes were stained with antibodies for protein targets where modulated cysteine residues were previously observed through proteomic analysis. GAPDH was used as a loading control.

### Methods

**DADPS vs. AZO Cleavable Biotin Linker Comparison**

*Cell Culture and Cellular Extract Preparation*

50 million K562 cells (ATCC source) were grown in IMDM (Gibco, 12440-053) supplemented with 9.1% FBS (Gibco, 16000-036) at 37 ˚C. Cells were pelleted by centrifugation in 50 mL conical tubes, washed with 10 mL PBS (Gibco, 10010-031), centrifuged at 700 x *g* for 3 minutes followed by vacuum aspiration and then frozen at -80 ˚C until further processing. The frozen cell pellet was thawed on ice and 1 mL of DPBS (Sigma-Aldrich, D8537-1L) supplemented with 1x EDTA-free protease/phosphatase inhibitor (Cell Signaling Technology, 5872) was added. Samples were lysed using a probe-tip sonication device (QSonica) and 4 cycles of 20 second sonication periods with 1 second intervals of sonication at 35% output. Samples were incubated on ice during sonication. Cellular extracts were clarified by centrifugation at 21000 x *g* and the supernatant (soluble fraction) was retained. Protein concentration was determined using a Bradford Assay.

*Probe Labeling and Bioorthogonal CuAAC Reaction for DADPS vs. Azobenzene Comparison*

1000 µg of cellular extract were aliquoted to a 2.0 mL Eppendorf tube and the total volume was adjusted to 500 µL. Samples were alkylated using 100 µM final concentration of IAAyne (Invitrogen, I10189). Following alkylation, the samples were subjected to copper-catalyzed azide-alkyne cycloaddition (CuAAC) reaction. 20 µL of a 44.2 mg/mL DMSO solution of Tris[(1-benzyl-1H-1,2,3-triazol-4-yl)methyl]amine (TBTA) (Sigma-Aldrich, 678937) was diluted in 180 µL of DMSO and further diluted to 1.7 mM using 800 µL of tert-Butanol (Sigma-Aldrich, 308250). 50 mM Tris(2-carboxyethyl)phosphine hydrochloride (TCEP) (Sigma-Aldrich, C4706) and 50 mM $CuSO_4$ (Sigma-Aldrich, 209198) were prepared in water (Sigma-Aldrich, W4502). 10 mM stock solutions of the dialkoxydiphenylsilane cleavable biotin linker (DADPS, Click Chemistry Tools, 1330) and azobenzene cleavable biotin linker (AZO, Click Chemistry Tools, 1040) were prepared in DMSO. A 62.5 µL master mix of CuAAC reagents was added to each protein solution for a final concentration of: 100 µM TBTA, 1 mM $CuSO_4$, 1.0 mM TCEP and 100 µM cleavable biotin-linker azide. Samples were incubated at room temperature in the dark for 60 minutes. Each cleavable linker was analyzed in three experimental replicates, with each experimental replicate originating from a different cell culture flask. Two sets of each cleavable biotin linker were processed, with one set being reserved for enrichment of denatured proteins and the other set being processed for enrichment of peptides post-trypsin digestion.

*Protein precipitation of CuAAC samples*

Following the CuAAC reaction, protein precipitation was performed using methanol-chloroform according to Wessel and Flügge[1]. Briefly, 500 µL of chilled methanol (EMD Millipore, MX0486) and 150 µL of chilled chloroform (Sigma-Aldrich, 366927) was added and the samples were vigorously vortexed. After 10 minutes of centrifugation at 21000 x *g* at 4 ˚C, the top aqueous layer was aspirated and 900 µL of chilled methanol was added. Samples were sonicated for one cycle of 20 seconds at 35% output, followed by centrifugation at 10 minutes of centrifugation at 21000 x *g* at 4 ˚C. The methanol supernatant was vacuum aspirated and the protein pellets were briefly air-dried.

*Streptavidin-biotin enrichment of denatured proteins post CuAAC reaction*

Samples designated for denatured protein enrichment were suspended in 650 µL of 2.5% SDS solution in DPBS, sonicated for one cycle of 20 seconds at 35% output, and heated at 65 ˚C for 5 minutes. Following centrifugation at 5000 x *g* for 5 minutes to pellet insoluble debris, the supernatant was transferred to a 15 mL conical tube. 100 µL of pre-washed streptavidin beads (Pierce Streptavidin Plus UltraLink, 53117) was washed with DPBS and added in 1 mL of DPBS to the conical tube. 8 mL of DPBS was added to dilute the final concentration of SDS to 0.2% and the mixture was incubated overnight on a rotating apparatus. The following day, the tubes were centrifuged at 700 x *g* for 3 minutes to pellet the beads. The supernatant was vacuum aspirated, and the beads were re-suspended in 1 mL of 1% SDS in DPBS and transferred to 1.5 mL Eppendorf tubes. Samples were washed twice with 1 mL of 1% SDS in DPBS, twice with 1 mL of 6M urea (Sigma-Aldrich, U5378) in DPBS, thrice with 1 mL DPBS and once with 1mL 100 mM ammonium bicarbonate pH 8.0 (Sigma-Aldrich, A6141), with all wash steps occurring by centrifuging the beads at 8000 x *g* for 1 min. 225 µL of ammonium bicarbonate pH 8.0 was added to the beads and 5 µL of 500 mM TCEP and 25 uL 500 mM CAA (VWR, 0898-100G) were added to reduce and alkylate free cysteines. Samples were incubated in a Thermomixer C (Eppendorf) for 30 minutes at 750 rpm at 37 ˚C. 1.0 µg of LysC (Wako, 125-05061) diluted in 10 µL of ammonium bircarbonate was added and the mixture was incubated for 1 hour at 37 ˚C at 1000 rpm in the Thermomixer. An aliquot of 1.0 µg of trypsin (Thermo Scientific, 90057) diluted in 10 µL of ammonium bicarbonate was added and the digest proceeded overnight. The next day the beads were centrifuged and the supernatant was aspirated and retained for further desalting. The beads were then washed 10 times with 1 mL of DPBS. DADPS-cleavable biotin linker samples were treated with 200 µL of 10% formic acid (Fisher, A117-50) and incubated for 20 minutes on a rotating apparatus. Azobenzene-cleavable biotin linker samples were treated with 200 µL of 25 mM sodium dithionite in DPBS (EMD Millipore, 1.06507) and incubated for 20 minutes on a rotating apparatus. All cleavages were repeated for a total of three times, and the individual eluent fractions for each respective cleavable biotin linker were combined. Sodium dithionite-cleaved peptides were acidified to 1% formic acid immediately prior to peptide desalting.

*Streptavidin-biotin enrichment of digested peptides post CuAAC reaction*

Samples designated for peptide enrichment were re-suspended in 300 µL of ammonium bicarbonate pH 8.0 containing 10 µg of LysC and incubated in a Thermomixer at 37 ˚C at 750 rpm for one hour. An additional 200 µL of ammonium bicarbonate was added containing 10 µg of trypsin and the digestion proceeded overnight. The next day, peptides were reduced and alkylated at a final concentration of 5 mM TCEP and 20 mM CAA for 30 minutes at 37 ˚C at 750 rpm in a Thermomixer. The peptide mixture was centrifuged at 5000 x *g* for 5 minutes to pellet any insoluble debris and 100 µL of streptavidin beads pre-washed in 100 mM ammonium bicarbonate pH 8.0 was added in 500 µL of ammonium bicarbonate to the supernatant of the digested peptides in a clean new tube. The tube was incubated on a rotating apparatus at room temperature overnight. The following day the beads were washed 10 times with 1 mL of DPBS. DADPS-cleavable biotin linker samples were treated with 200 µL of 10% formic acid and incubated for 20 minutes on a rotating apparatus. Azobenzene-cleavable biotin linker samples were treated with 200 µL of 25 mM sodium dithionite in DPBS and incubated for 20 minutes on a rotating apparatus. All cleavages were repeated for a total of three times, and the individual eluent fractions for each respective cleavable biotin linker were combined. Sodium dithionite-cleaved peptides were acidified to 1% formic acid immediately prior to peptide desalting.

*Peptide desalting and fractionation*

Peptides from chemical cleavage and on-bead digestion were desalted using styrene-divinylbenzene reversed phase sulfonate StageTips (3M Empore, 2241) according to Kulak *et al* [2]. Briefly, 6 layers of material was plugged in a 200 µL pipette tip, the pipette tip was conditioned with 100 µL of the following solvents: methanol, 80% MeCN 0.1% FA, and 0.2% TFA. Acidified samples were loaded onto the StageTips and washed with 100 µL of the following solvents: 1% TFA in isopropanol, 0.2% TFA. StateTips were then transferred to new tubes and cleaved peptides were fractionated by eluting with 100 µL of the following solvent into a new tube for each fraction: 100 mM ammonium formate in 40% MeCN 0.5% FA (fraction 1), 150 mM ammonium formate in 60% MeCN 0.5% FA (fraction 2), and 80% MeCN 5% $NH_4OH$ (fraction 3). Peptides resulting from on-bead digestion were not fractionated and directly eluted from the StageTips with 100 µL 80% MeCN 5% $NH_4OH$. Samples were vacuum centrifuged in a Speedvac at 45 °C until dryness and reconstituted in 35 µL of 1% FA.

*rGSTO1 analysis for sulfation*

10 µg of recombinant GSTO1 (Sigma-Aldrich, GS75) was aliquoted to 1.5 mL Eppendorf tubes and the total volume was adjusted to 45 µL. Samples were alkylated using 100 µM final concentration of IAAyne. Following alkylation, the samples were subjected to copper-catalyzed azide-alkyne cycloaddition (CuAAC) reaction as previously described. A 5 µL master mix of CuAAC reagents was added to each protein solution for a final concentration of: 100 µM TBTA, 1 mM $CuSO_4$, 1.0 mM TCEP and 100 µM cleavable AZO biotin-linker azide. Samples were incubated at room temperature in the dark for 60 minutes. Samples were diluted to a final volume of 100 µL by adding 50 µL of 50 mM sodium dithionite in DPBS. Samples were incubated at room temperature in the dark for 30 minutes. One sample was acidified to 1% FA while the control sample had an equivalent volume of water added to it. Samples were then precipitated using chloroform-methanol as previously described and digested using LysC (1:10) and trypsin (1:10) overnight in a Thermomixer C with shaking. Following digestion, samples were reduced and alkylated with additional TCEP (5 mM) and CAA (20 mM). Samples were desalted using SDB-RPS Stage Tips as previously described and reconstituted in 40 µL of 1% FA.

*LC-MS/MS analysis of peptides*

Samples were analyzed using a Thermo Scientific Orbitrap Fusion tribrid MS instrument. 10 µL of peptide samples were injected using a Thermo Scientific EASY-nLC 1200 onto a 75 µm x 2 cm 3 µm Acclaim PepMap 100 C18 trapping column (Thermo, 164946) followed by separation using a 75 µm x 25 cm 2 µm, 100A EasySpray PepMap RSLC C18 analytical column (Thermo, ES802). Peptide samples from on-bead digests (1 µL) were analyzed using a Thermo Scientific Ultimate 3000 LC. 0.1% formic acid in water (Optima LC-MS grade) mobile phase A, and 0.1% formic acid in 80% MeCN (Optima LC-MS grade) mobile phase B were utilized as solvents. Peptides were separated using a linear gradient from 5% to 35% B over 46 minutes at a flow rate of 250 nL/min. The Orbitrap Fusion MS was operated in data-dependent mode in positive ion mode. MS1 full scan was performed at 60000 resolution in the Orbitrap scanning from 300-1500 *m/z*. AGC target setting was 4.0e5 and maximum IT was 50 ms. MIPS was set to Peptide, Intensity Threshold was set to 2.0e4, charge state 2-7 were selected and dynamic exclusion was set to 30 seconds with n exclusion after 1 and a high/low mass tolerance of 10 ppm. For HCD analysis of peptides, data-dependent MS2 in the Orbitrap was performed at 15000 resolution with a fixed first mass of 120 *m/z*. Quadrupole isolation was enabled with an isolation window of 1.6 *m/*z, HCD activation enabled and collision energy set to 28%. AGC target setting was 5.0e4 and maximum IT was 110 ms. For CID analysis of peptides, data-dependent MS2 in Linear Ion Trap was performed using Turbo scan rate selected. Quadrupole isolation was enabled with an isolation window of 1.6 *m/z*, CID activation set to a collision energy of 35%, with activation time of 10 ms and Q of 0.25. AGC target setting was 7.5e3 and maximum IT was 75 ms. A Top Speed cycle time of 3 seconds was utilized and lock mass for 445.12 *m/z* was enabled for all analyses.

*Bioinformatic Analysis of MS Data*

FragPipe (v9.1) was utilized with MSFragger (v20190222)[3] and philosopher (v201903191601) to perform mass tolerant database searching of samples acquired using HCD and Orbitrap MS2 detection. Raw files were searched against a canonical sequence database of the Swissprot-validated human proteome that was downloaded from Uniprot on September 6$^{th}$, 2017 (20320 entries) and appended with the cRAP protein sequences from the Global Proteome Machine (https://www.thegpm.org/crap/). A reverse decoy database was included. Trypsin was selected with 2 enzyme termini and 1 missed cleavage. Default open search parameters were enabled with the following alterations: precursor mass tolerance 0-500 Da, precursor true tolerance 20 ppm, variable modification on M (+15.99490) and fixed modification on C (+57.021464). PeptideProphet[4] was enabled with the following settings: --nonparam --expectscore --decoyprobs --masswidth 1000.0 --clevel -2. Crystal-C was enabled with the following settings: max charge 6, mass tolerance 20 ppm, number of isotopes 3, precursor isolation window 1.6. Localization of mass modifications was performed with PTMProphetParser.exe (TPP v5.1.0) using the following parameters: VERBOSE MZTOL 0.02 MASSDIFFMODE M:15.9949, C57.021464. The Trans-Proteomic Pipeline (v5.1.0)[5] was used to inspect spectra.

Database searching of raw files acquired using CID and LTQ MS2 detection was performed using MaxQuant (v1.6.5.0)[6, 7] in order to tabulate the number of unique cysteine-sites identified using each cleavable linker. Standard label-free searching was selected. Andromeda search engine was used to perform a database search created from a canonical sequence database of the Swissprot-validated human proteome that was downloaded from Uniprot on September 6$^{th}$, 2017 (20320 entries). Trypsin/P with two missed cleavages was selected. Minimum peptide length was set to 7. Carbamidomethyl C (+57.021463 Da) was set as a fixed modification. Variable modifications used were: Deamidation N/Q (+0.984015 Da), Oxidation M (+15.994914 Da), DADPS-residual C (+181.121512 Da), AZO-residual C(+216.101111) and AZO-SULFO-residual C (+296.057925). PSM and Protein FDR were set to 1%. The remainder of the search parameters were left at default values. MaxQuant output results were loaded into Perseus (v1.6.5.0)[8] and reverse hits and contaminants were filtered. Sites with a localization score greater than 0.80 were retained for analyses.

For recombinant GSTO1 samples, FragPipe(v9.1) FragPipe (v9.1) was utilized with MSFragger (v20190222) and philosopher (v201903191601) to perform standard database searching. Files were searched against a Swissprot-validated human proteome as previously described. Trypsin was selected with 2 enzyme termini and 1 missed cleavage. Default closed search parameters were enabled with the following alterations: precursor mass tolerance 0-20 ppm, precursor true tolerance 20 ppm, isotope error 0/1/2, variable modification on M (+15.99490), C (+216.1011), C(+296.0579) and fixed modification on C (+57.021464). PeptideProphet[4] was enabled with the following settings: --nonparam --expectscore --decoyprobs --accmass --ppm. Extracted ion chromatograms of modified peptides were performed using Thermo Xcalibur Qual Browser (v3.0.63).

**<u>Profiling Dacomitinib Using DADPS Cleavable Biotin Azide Linker</u>**

*Cell Culture and Cellular Extract Preparation*

25 million A431 cells (ATCC source) were grown in DMEM (Gibco, 11965-092) supplemented with 9.1% FBS (Gibco, 16000-036) at 37 ˚C in 15 cm cell culture dishes. Cells were treated with 5 µM dacomitinib (Sigma-Aldrich, PZ0330) or DMSO in fresh media supplemented with FBS for 2 hours at 37 ˚C. Treatments were performed in triplicate and each replicate was performed on a different day. Upon completion of treatment, media was vacuum aspirated and cells were harvested by placing the cell culture dishes on a bed of ice and scraping off the cells and re-suspending in chilled PBS. Cells were pelleted by centrifugation in 50 mL conical tubes, washed further with 10 mL PBS (Gibco, 10010-031), centrifuged at 700 x *g* for 3 minutes followed by vacuum aspiration and then frozen at -80 ˚C until further processing. The frozen cell pellet was thawed on ice and 1 mL of DPBS (Sigma-Aldrich, D8537-1L) supplemented with 1x EDTA-free protease/phosphatase inhibitor (Cell Signaling Technology, 5872) was added. Samples were lysed using a Bioruptor Plus device (Diagenode, B01020001 UCD-300 TM) at 4 ˚C, running 10 cycles of 15 second sonication periods on HIGH setting followed by 45 seconds of resting periods. 1 µL of benzonase nuclease (Sigma-Aldrich, E1014) was added and cellular extracts were incubated on ice for 30 minutes. Cellular extracts were then clarified by centrifugation at 21000 x *g* and the supernatant was retained. Protein concentration was determined using a Bradford Assay.

*SDS-PAGE In-Gel Fluorescence To Confirm Dacomitinib Treatment*

50 µg of protein from each replicate was incubated with 1 µM dacomitinib probe in 45 µL at room temperature in the dark. Following incubation, the samples were subjected to copper-catalyzed azide-alkyne cycloaddition (CuAAC) reaction. 20 µL of a 44.2 mg/mL DMSO solution of Tris[(1-benzyl-1H-1,2,3-triazol-4-yl)methyl]amine (TBTA) (Sigma-Aldrich, 678937) was diluted in 180 µL of DMSO and further diluted to 1.7 mM using 800 µL of tert-Butanol (Sigma-Aldrich, 308250). 50 mM Tris(2-carboxyethyl)phosphine hydrochloride (TCEP) (Sigma-Aldrich, C4706) and 50 mM $CuSO_4$ (Sigma-Aldrich, 209198) were prepared in water (Sigma-Aldrich, W4502). 0.5 mM Tetramethylrhodamine 5-

Carboxamido-(6Azidohexanyl)(TAMRA)-azide (Invitrogen, T10182) was prepared in DMSO. A 5.5 µL master mix of CuAAC reagents was added to each protein solution for a final concentration of: 100 µM TBTA, 1 mM $CuSO_4$, 0.5 mM TCEP and 10 µM TAMRA-azide(T10182). Samples were incubated at room temperature in the dark for 60 minutes. Following CuAAC reaction, 15 µL of 4x LDS sample buffer (Invitrogen, NP0007) and 5 µL of 10x reducing agent (Invitrogen, NP0009) was added to each sample and incubated at 70 ˚C for 10 minutes. Samples were separated by SDS-PAGE using a 4-12% NuPAGE Bis-Tris gel (Invitrogen, NP0335) for 90 minutes at 150V using MES buffer (Invitrogen, NP0002). Following separation, the gel was fixed using 50% methanol (Sigma-Aldrich, 34860) and 7% acetic acid (Sigma-Aldrich, 695092) for 30 minutes followed by incubation in 40% methanol overnight, and another incubation for 30 minutes in destain solution (10% methanol, 7 % acetic acid). For total stain, gels were incubated in SYPRO (Invitrogen, S12000) and incubated in destain solution overnight. Gels were imaged using a BioRad ChemiDoc MP imager with green epi-illumination for excitation and a 605/50 filter for TAMRA and UV trans-illumination for excitation and a 605/50 filter for SYPRO. Following in-gel fluorescence image capture, total protein stain was performed using SYPRO and subsequently imaged.

*Western Blot Analysis*

20 µg of sample was used for probing with EGFR antibodies prior to processing samples for mass spectrometry analysis. 10 µg of sample was used for subsequent analyses. 4x LDS sample buffer (Invitrogen, NP0007) and 10x reducing agent (Invitrogen, NP0009) was added to each sample and incubated at 70 ˚C for 10 minutes. Samples were separated by SDS-PAGE using a 4-12% NuPAGE Bis-Tris gel (Invitrogen, NP0335) for 90 minutes at 150V using MES buffer (Invitrogen, NP0002). Following separation, gels were transferred to distilled water prior to membrane transfer using an iBlot2 system (Invitrogen). Gels were transferred to PVDF membrane (Invitrogen, IB24001) using the template P0 transfer method. Immediately following transfer, membranes were incubated in light-impenetrable boxes (LI-COR) with TBS or PBS buffer. Membranes were blocked using LI-COR Oddyssey Blocking Buffer in TBS (927-50000) or PBS (927-40000) for one hour at room temperature. Primary antibodies were used 1:1000 and incubated overnight in 50% LI-COR Odyssey Blocking Buffer with 0.1% Tween-20 additive in either TBS or PBS. Antibodies used: ELF3 (MAB5787, R&D Systems), EGFR pY1068 (2234S, Cell Signaling Technologies), EGFR (ab52894, abcam), GSK3B (12456S, Cell Signaling Technologies), GSK3B pS9 (9336S, Cell Signaling Technologies), GSK3B pY216 (ab75745, abcam), FRA1 (FOSL1) (5281S, Cell Signaling Technologies), NUR77 (NR4A1) (ab109180, abcam), GAPDH (MAB374, EMD Millipore). Membranes were washed the next day in TBS or PBS with 0.1% Tween-20 additive followed by incubation with secondary antibodies for 30 minutes at 1:10000 in 50% LI-COR Odyssey Blocking Buffer with 0.1% Tween-20 additive in either TBS or PBS at room temperature. Secondary antibodies used: IRDye Goat-anti-Mouse 800CW (926-32210, LI-COR), IRDye Goat-anti-Rabbit 800CW (926-32211, LI-COR), IRDye Goat-anti-Mouse 680RD (926-68070, LI-COR). Membranes were then washed in TBS or PBS with 0.1% Tween-20 additive followed by rinse in TBS or PBS prior to imaging using an Odyssey CLx scanner (LI-COR).

*Multiplexed Profiling of Dacomitinib Against Cysteinome Using IAAyne and DADPS Biotin Azide*

500 µg of cellular extract were aliquoted to a 2.0 mL Eppendorf tube and the total volume was adjusted to 500 µL. Samples were alkylated using 100 µM final concentration of IAAyne (Invitrogen, I10189) for

60 minutes at room temperature in the dark. Following labeling, the samples were subjected to copper-catalyzed azide-alkyne cycloaddition (CuAAC) reaction. 20 µL of a 44.2 mg/mL DMSO solution of Tris[(1-benzyl-1H-1,2,3-triazol-4-yl)methyl]amine (TBTA) (Sigma-Aldrich, 678937) was diluted in 180 µL of DMSO and further diluted to 1.7 mM using 800 µL of tert-Butanol (Sigma-Aldrich, 308250). 50 mM Tris(2-carboxyethyl)phosphine hydrochloride (TCEP) (Sigma-Aldrich, C4706) and 50 mM $CuSO_4$ (Sigma-Aldrich, 209198) were prepared in water (Sigma-Aldrich, W4502). 10 mM stock solution of the dialkoxydiphenylsilane cleavable biotin linker (DADPS, Click Chemistry Tools, 1330) was prepared in DMSO. A 62.5 µL master mix of CuAAC reagents was added to each protein solution for a final concentration of: 100 µM TBTA, 1 mM $CuSO_4$, 1.0 mM TCEP and 100 µM cleavable biotin-linker azide. Samples were incubated at room temperature in the dark for 60 minutes. Immediately following the CuAAC reaction, 5 µL of 500 mM TCEP and 20 µL of 500 mM CAA was added and incubated for 20 minutes at room temperature in the dark. For multiplexed profiling, the samples were then processed using an adapted protocol from Navarrete-Perea *et al*[9] for TMT labeling in combination with the previously described protocol for the DADPS cleavable biotin azide linker. Protein precipitation was performed using methanol-chloroform as previously described. After methanol washing, methanol was vacuum aspirated and 100 µL of 200 mM 3-[4-(2-hydroxyethyl)piperazin-1-yl]propane-1-sulfonic acid (EPPS) pH 8.5 containing 10 µg of LysC was added. Samples were incubated in a Thermomixer C with shaking at 37 ˚C at 800 rpm for one hour. Another 100 µL of 200 mM EPPS pH 8.5 containing 10 µg of trypsin was added and the digestion proceeded overnight. The next morning the digest was centrifuged at 5000 x *g* for 5 minutes to pellet any insoluble debris. Supernatant was retained and transferred to a new tube. TMT 6-plex reagent (Thermo Scientific, 90066) was thawed to room temperature, and 0.8 mg of reagent was re-suspended in 87 µL of MeCN which was then vortexed and added to each replicate. Samples were incubated for 2 hours at 25 ˚C in a Thermomixer C with shaking at 800 rpm. The reaction was quenched by adding 1.7 µL of 50 % hydroxylamine solution (EMD Millipore, 8.14441.0100). Sample volume was reduced to ~ 150 µL and an aliquot from each replicate was mixed 1:1:1:1:1:1 to perform a ratio check. Samples were desalted for unfractionated analysis as previously described. Using normalization factors calculated by the ratio check, TMT-labeled samples were mixed together into a new 2.0 mL tube. An aliquot of this mixture was retained to serve as "input" for total proteome quantitation. 200 µL of pre-washed streptavidin resin (Pierce Streptavidin Plus UltraLink, 53117) was re-suspended in 1 mL of DPBS and added to the TMT-labeled peptides. The tube was incubated on a rotating apparatus at room temperature overnight. The following day the beads were washed 10 times with 1 mL of DPBS. Samples were treated with 200 µL of 10% formic acid in 5% MeCN and incubated for 20 minutes on a rotating apparatus, repeated for a total of three times, and the individual eluent fractions for each respective cleavable biotin linker were combined. Samples were fractionated and desalted as previously described. Samples were vacuum centrifuged in a Speedvac at 45 ˚C until dryness and reconstituted in 25 µL of 1% FA.

*LC-MS/MS Analysis of Peptides*

Samples were analyzed using a Thermo Scientific Orbitrap Fusion Lumos tribrid MS instrument. 7.5 µL of peptide samples were injected using a Thermo Scientific EASY-nLC 1200 onto a 75 µm x 2 cm 3 µm Acclaim PepMap 100 C18 trapping column (Thermo, 164946) followed by separation using a 75 µm x 25 cm 2 µm, 100A EasySpray PepMap RSLC C18 analytical column (Thermo, ES802). 0.1% formic acid in water (Optima LC-MS grade) mobile phase A, and 0.1% formic acid in 80% MeCN (Optima LC-MS grade) mobile phase B were utilized as solvents. Peptides were separated using a linear gradient from 5% to 35% B over 106 minutes at a flow rate of 250 nL/min. The Orbitrap Fusion Lumos MS was operated in data-dependent mode in positive ion mode using a SPS-MS3 method. MS1 full scan was performed at 120000 resolution in the Orbitrap scanning from 375-1800 *m/z*. AGC target setting was 4.0e5 and maximum IT was 50 ms. MIPS was set to Peptide, Intensity Threshold was set to 5.0e3, charge state 2-7 were selected and dynamic exclusion was set to 60 seconds with n exclusion after 1 and a high/low mass tolerance of 10 ppm. Quadrupole isolation was enabled with an isolation window of 0.7 *m/z*, CID activation set to a collision energy of 35%, with activation time of 10 ms and Q of 0.25. AGC target setting was 1.0e4 and maximum IT was 50 ms. Following each MS2 acquisition, a synchronous precursor scan (SPS)-MS3 scan was performed with 10 SPS Precursors, HCD collision energy set to 65%, an AGC target of 1.0e5 and a maximum IT of 105 ms. A Top Speed cycle time of 3 seconds was utilized and lock mass for 445.12 *m/z* was enabled for all analyses.

*Bioinformatic Analysis of LC-MS/MS Data*

Database searching of raw files was performed using MaxQuant (v1.6.5.0)[6, 7]. Reporter ion MS3 with isotopic correction factor adjustment was used. Andromeda search engine was used to perform a database search created from a canonical sequence database of the Swissprot-validated human proteome that was downloaded from Uniprot on September 6$^{th}$, 2017 (20320 entries).Trypsin/P with two missed cleavages was selected. Minimum peptide length was set to 7. Carbamidomethyl C (+57.021463 Da) was set as a fixed modification. Variable modifications used were: Deamidation N/Q (+0.984015 Da), Oxidation M (+15.994914 Da) and DADPS-residual C (+181.121512 Da). PSM and Protein FDR were set to 1%. The remainder of the search parameters were left at default values. For database searching of the peptides used as input for enrichment to quantify the proteome, only unmodified peptides (unique + razor) were used for quantification. MaxQuant output results were loaded into Perseus (v1.6.5.0)[8] and reverse hits and contaminants were filtered out. Files were exported as tab-separated text files and imported into RStudio (v1.1.453, R v3.5.3). edgeR (v3.24.3) was used to normalize intensities across TMT channels using trimmed means of M values[10]. Intensities were then log2 transformed and ratios between DMSO/dacomitinib treatment were calculated by subtracting the mean log2 intensities of dacomitinib-treated samples from the mean log2 intensities of DMSO-treated samples. Analyses were visualized and plotted using ggplot2 (v3.1.1).

**Dacomitinib Probe Synthesis**

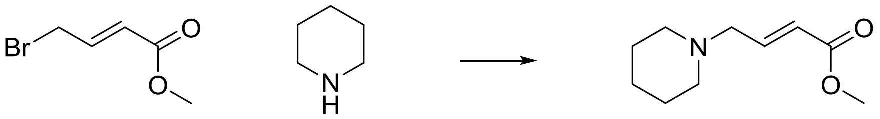

In a 100 mL round bottom flask was added (E)-methyl 4-bromobut-2-enoate (5 g, 27.9 mmol) in CH2Cl2 (30 ml) to give a colorless solution. piperidine (5.52 ml, 55.9 mmol) in CH2Cl2 (5 ml) was added dropwise and the reaction was stirred at RT for 2 hrs at which point a slurry formed. The reaction was concentrated in vacuo and purified via silica gel chromatography (0-40% EtOAc in heptanes gradient). The title compound was isloated as a yellow oil (4.24 g, 83% yield). 1H NMR (400 MHz, Chloroform-d) δ 6.99 (dt, J = 15.7, 6.2 Hz, 1H), 5.97 (dt, J = 15.7, 1.6 Hz, 1H), 3.73 (s, 3H), 3.10 (dd, J = 6.3, 1.7 Hz, 2H), 2.39 (t, J = 5.4 Hz, 4H), 1.64 - 1.54 (m, 4H), 1.49 - 1.38 (m, 2H).

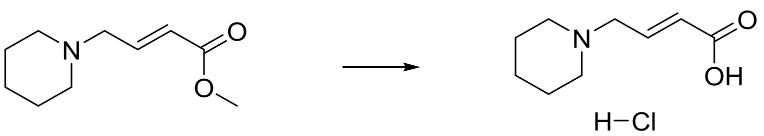

In a 200 mL round bottom flask was added (E)-methyl 4-(piperidin-1-yl)but-2-enoate (4.2 g, 22.92 mmol) in Dioxane (57.3 ml) to give a colorless solution. hydrogen chloride (2 M in H2O, 22.92 ml, 45.8 mmol) was added and heated to 80 °C over the weekend. Solvent was removed in vacuo and the resulting oil was azeotroped with toluene. A white solid formed which was washed with 50 mL MTBE, filtered and washed with EtOAc. Dried in vacuum oven overnight. 1H NMR (501 MHz, DMSO-d6) δ 10.88 (s, 1H), 6.86 (dt, J = 15.7, 7.1 Hz, 1H), 6.14 (dd, J = 15.6, 1.5 Hz, 1H), 3.83 (dd, J = 7.1, 1.4 Hz, 2H), 2.83 - 2.79 (m, 4H), 1.80 - 1.55 (m, 5H), 1.37 - 1.33 (m, 1H). 13C NMR (101 MHz, DMSO-d6) d 166.34, 136.16, 129.71, 56.01, 52.22, 22.65, 21.63. MS(ESI) m/z 170.3 [M+H].

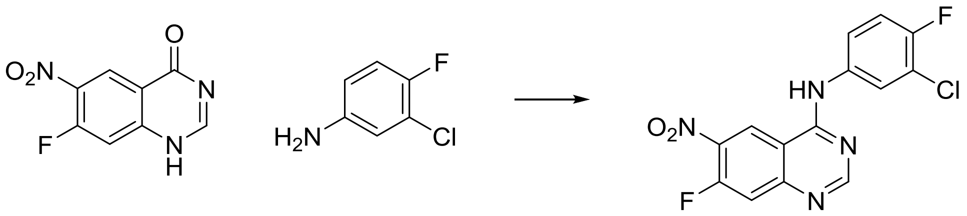

In a 200 mL round bottom flask was added 7-fluoro-6-nitroquinazolin-4(1H)-one (4 g, 19.13 mmol) and sulfurous dichloride (75 ml, 1028 mmol) to give a yellow suspension. Three drops of DMF were added. The mixture was heated to 85 °C for 6 hrs, at which point the reaction mixture became a yellow solution. The thionyl chloride was removed in vacuo to yield an off-white solid.

The residue was suspended in CH2Cl2 (75 ml). 3-chloro-4-fluoroaniline (2.92 g, 20.08 mmol) in Ethanol (EtOH) (20 ml) was added and the reaction stirred at RT. After 5-10 minutes, the reaction mixture became a thick yellow slurry. 100 mL heptanes was added and the reaction filtered. The solid was washed with MTBE and dried in vacuo. Residue was dry loaded onto silica and the compound was purified via normal phase chromatography (0-100% EtOAc in heptanes) to yield the title compound (3.25 g, 51% yield). 1H NMR (400 MHz, DMSO-d6) δ 10.49 (s, 1H), 9.55 (d, J = 7.9 Hz, 1H), 8.70 (s, 1H), 8.14 - 8.07 (m, 1H), 7.82 (d, J = 12.5 Hz, 1H), 7.76 (dd, J = 8.1, 4.4 Hz, 1H), 7.46 (t, J = 9.1 Hz, 1H). 13C NMR (101 MHz, DMSO-d6) δ 158.39, 158.26, 157.78, 155.18 (d, J = 7.1 Hz), 154.06 (d, J = 13.5 Hz), 152.79, 135.60, 135.48 (d, J = 3.3 Hz), 124.44 (d, J = 4.7 Hz), 123.22 (d, J = 7.1 Hz), 119.02 (d, J = 18.5 Hz), 116.77 (d, J = 21.8 Hz), 115.08 (d, J = 20.2 Hz), 111.29 (d, J = 1.6 Hz). MS(ESI) m/z 337.2 [M+H].

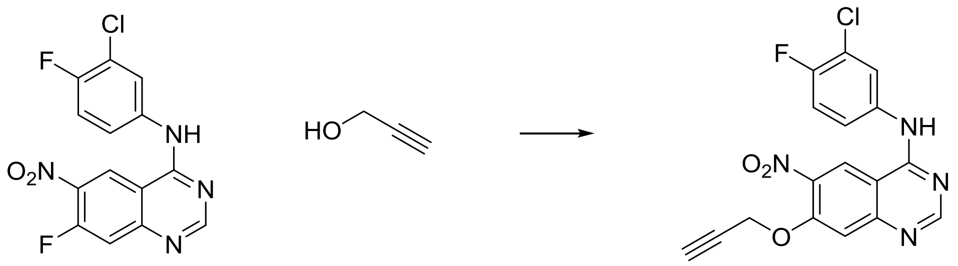

In a 100 mL round-bottomed flask was added sodium hydride (0.356 g, 8.91 mmol) in Dimethyl formamide (DMF) (20 ml) to give a white suspension. This was cooled to 0 °C. prop-2-yn-1-ol (0.531 ml, 9.21 mmol) was added in Dimethyl formamide (DMF) (5 ml) and the reaction was stirred at 0 °C for 30 minutes. N-(3-chloro-4-fluorophenyl)-7-fluoro-6-nitroquinazolin-4-amine (1 g, 2.97 mmol) in Dimethyl formamide (DMF) (10 ml) was added dropwise and the reaction stirred at 0 °C for 2 hrs, at which point the reaction was deemed complete by LC.

The reaction was poured into 150 mL DCM, and 200 mL 1 M HCl was added. The aqueous layer was extracted with 150 mL DCM two times. Extracts were washed with brine, concentrated and dried in vacuo to yield the title compound (1.05 g, 95% yield). 1H NMR (400 MHz, DMSO-d6) δ 10.24 (s, 1H), 9.27 (s, 1H), 8.64 (s, 1H), 8.14 (dd, J = 6.9, 2.6 Hz, 1H), 7.79 (ddd, J = 9.0, 4.3, 2.6 Hz, 1H), 7.53 (s, 1H), 7.43 (t, J = 9.1 Hz, 1H), 5.18 (d, J = 2.4 Hz, 2H), 3.77 (d, J = 2.2 Hz, 1H). 13C NMR (101 MHz, DMSO-d6) δ 157.85, 157.54, 154.76, 153.11, 152.34, 152.24, 138.74, 136.54, 123.88, 122.70 (d, J = 6.9 Hz), 122.17, 118.87 (d, J = 18.4 Hz), 116.58 (d, J = 21.7 Hz), 110.96, 108.83, 79.88, 77.70, 57.45. MS(ESI) m/z 373.2 [M+H].

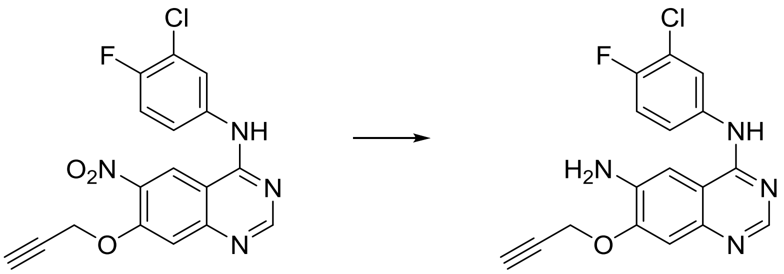

In a 20 mL round-bottomed flask was N-(3-chloro-4-fluorophenyl)-6-nitro-7-(prop-2-yn-1-yloxy)quinazolin-4-amine (610 mg, 1.637 mmol), ammonia hydrochloride (525 mg, 9.82 mmol), and zinc (535 mg, 8.18 mmol) in Methanol (MeOH) (20 ml) to give a yellow suspension. The reaction mixture was sonicated for 2 minutes with swirling, then stirred overnight at RT.

The sample was purified via reverse phase column chromatography to yield the final compound as an orange solid (450 mg, 80% yield). 1H NMR (400 MHz, DMSO-d6) d 9.45 (s, 1H), 8.38 (s, 1H), 8.20 (dd, J = 6.9, 2.6 Hz, 1H), 7.82 (ddd, J = 9.1, 4.4, 2.7 Hz, 1H), 7.46 (s, 1H), 7.39 (t, J = 9.1 Hz, 1H), 7.22 (s, 1H), 5.37 (s, 2H), 5.04 (d, J = 2.4 Hz, 2H), 3.68 (t, J = 2.3 Hz, 1H). 13C NMR (101 MHz, DMSO-d6) d 155.53, 154.33, 151.92, 150.80 (d, J = 2.4 Hz), 144.83, 138.99, 137.93 (d, J = 3.0 Hz), 122.95, 121.91 (d, J = 6.7 Hz), 119.05 (d, J = 18.2 Hz), 116.83 (d, J = 21.5 Hz), 111.28, 107.96, 101.84, 79.42, 79.12, 56.49. MS(ESI) m/z 343.2 [M+H].

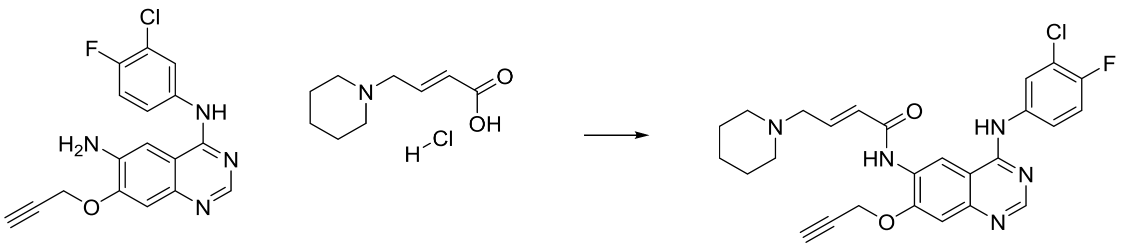

In a 4 mL vial was (E)-4-(piperidin-1-yl)but-2-enoic acid hydrochloride (75 mg, 0.365 mmol) in Dichloromethane (DCM) (2 ml) to give a white suspension. The mixture was cooled to 0 °C. oxalyl dichloride (0.048 ml, 0.547 mmol) was added, followed by 1 drop of DMF. The reaction was capped and stirred at RT for 1 hr, at which point it was allowed to warm to RT. When the reaction became clear, an LCMS was taken (MeOH quench) that verified complete conversion to the acid chloride. This was used directly in the next step without modification.

In a 20 mL vial was N4-(3-chloro-4-fluorophenyl)-7-(prop-2-yn-1-yloxy)quinazoline-4,6-diamine (62.5 mg, 0.182 mmol) in Tetrahydrofuran (THF) (10 ml) to give a brown solution. N-ethyl-N-isopropylpropan-2-amine (0.318 ml, 1.823 mmol) was added and the mixture cooled to 0 °C. The stock solution of acid chloride was added dropwise. The reaction was warmed to RT and allowed to stir for 3 hrs at which point the reaction appeared to stall. The material was directly purified via reverse phase HPLC/MS and concentrated to yield the final compound (52 mg, 58% yield). 1H NMR (400 MHz, DMSO-d6) δ 9.81 (s, 1H), 8.99 (s, 1H), 8.62 (s, 1H), 8.21 (dd, J = 6.9, 2.6 Hz, 1H), 7.89 (ddd, J = 9.1, 4.4, 2.7 Hz,

1H), 7.56 - 7.45 (m, 2H), 6.90 (dt, J = 15.4, 6.1 Hz, 1H), 6.64 (d, J = 15.4 Hz, 1H), 5.18 (d, J = 2.4 Hz, 2H), 3.78 (t, J = 2.3 Hz, 1H), 3.19 (dd, J = 6.1, 1.6 Hz, 2H), 2.45 (t, J = 5.3 Hz, 4H), 1.61 (p, J = 5.5 Hz, 4H), 1.51 - 1.43 (m, 2H). 13C NMR (101 MHz, DMSO-d6) δ 172.83, 163.81, 156.89, 154.52, 154.05, 153.33, 152.11, 148.65, 142.18, 136.83 (d, J = 3.0 Hz), 127.35, 125.85, 123.68, 122.53 (d, J = 6.8 Hz), 118.81 (d, J = 18.3 Hz), 116.77, 116.54 (d, J = 21.5 Hz), 109.39, 108.30, 79.42, 78.45, 59.38, 56.48, 54.15, 25.57, 23.92. MS(ESI) m/z 494.3 [M+H].

### References

1. Wessel, D., and Flügge, U. I. (1984) A method for the quantitative recovery of protein in dilute solution in the presence of detergents and lipids, *Analytical Biochemistry 138*, 141-143.
2. Kulak, N. A., Pichler, G., Paron, I., Nagaraj, N., and Mann, M. (2014) Minimal, encapsulated proteomic-sample processing applied to copy-number estimation in eukaryotic cells, *Nat Methods 11*, 319-324.
3. Kong, A. T., Leprevost, F. V., Avtonomov, D. M., Mellacheruvu, D., and Nesvizhskii, A. I. (2017) MSFragger: ultrafast and comprehensive peptide identification in mass spectrometry-based proteomics, *Nat Methods 14*, 513-520.
4. Keller, A., Nesvizhskii, A. I., Kolker, E., and Aebersold, R. (2002) Empirical Statistical Model To Estimate the Accuracy of Peptide Identifications Made by MS/MS and Database Search, *Analytical Chemistry 74*, 5383-5392.
5. Deutsch, E. W., Mendoza, L., Shteynberg, D., Slagel, J., Sun, Z., and Moritz, R. L. (2015) Trans-Proteomic Pipeline, a standardized data processing pipeline for large-scale reproducible proteomics informatics, *Proteomics Clin Appl 9*, 745-754.
6. Cox, J., and Mann, M. (2008) MaxQuant enables high peptide identification rates, individualized p.p.b.-range mass accuracies and proteome-wide protein quantification, *Nat Biotechnol 26*, 1367-1372.
7. Tyanova, S., Temu, T., and Cox, J. (2016) The MaxQuant computational platform for mass spectrometry-based shotgun proteomics, *Nat Protoc 11*, 2301-2319.
8. Tyanova, S., Temu, T., Sinitcyn, P., Carlson, A., Hein, M. Y., Geiger, T., Mann, M., and Cox, J. (2016) The Perseus computational platform for comprehensive analysis of (prote)omics data, *Nat Methods 13*, 731-740.
9. Navarrete-Perea, J., Yu, Q., Gygi, S. P., and Paulo, J. A. (2018) Streamlined Tandem Mass Tag (SL-TMT) Protocol: An Efficient Strategy for Quantitative (Phospho)proteome Profiling Using Tandem Mass Tag-Synchronous Precursor Selection-MS3, *J Proteome Res 17*, 2226-2236.
10. Robinson, M. D., and Oshlack, A. (2010) A scaling normalization method for differential expression analysis of RNA-seq data, *Genome Biol 11*, R25.
